# Robust Prediction of Patient-Specific Clinical Response to Unseen Drugs From *in vitro* Screens Using Context-aware Deconfounding Autoencoder

**DOI:** 10.1101/2021.05.20.445055

**Authors:** Di He, Qiao Liu, Lei Xie

## Abstract

Accurate and robust prediction of patient-specific responses to drug treatments is critical for drug development and personalized medicine. However, patient data are often too scarce to train a generalized machine learning model. Although many methods have been developed to utilize cell line data, few of them can reliably predict individual patient clinical responses to new drugs due to data distribution shift and confounding factors. We develop a novel Context-aware Deconfounding Autoencoder (CODE-AE) that can extract common biological signals masked by context-specific patterns and confounding factors. Extensive studies demonstrate that CODE-AE effectively alleviates the out-of-distribution problem for the model generalization, significantly improves accuracy and robustness over state-of-the-art methods in both predicting patient-specific *ex vivo* and *in vivo* drug responses purely from *in vitro* screens and disentangling intrinsic biological signals from confounding factors. Using CODE-AE, we screened 50 drugs for 9,808 cancer patients and discovered novel personalized anti-cancer therapies and drug-response biomarkers.

## 1 Introduction

Omics profiling, particularly transcriptomics, is a powerful technique to characterize cellular activity under various conditions, allowing developing machine learning models for personalized phenotype drug screening [1, 2, 3]. However, the success of such predictive models largely relies on the availability of sufficient amounts of data with coherent and comprehensive annotations. In clinical, we are often short of a large number of coherent *in vivo* patient data with drug treatment and response history. As a result, most drug response predictive studies to date have mainly utilized transcriptomic profiles from panels of *in vitro* cancer cell lines. Although such an approach is promising, the utility of drug response predictive models built with *in vitro* data is often limited when applied to actual patients due to the genetic and environmental differences between *in vitro* cell lines and patient-derived tissue samples as well as various confounding factors and overwhelming context-specific patterns that may mask intrinsic biological signals.

The inability to predict patient-specific *in vivo* drug responses from *in vitro* screening data using a machine learning approach originates from a fundamental challenge of out-of-distribution (OOD) problem. The underlying assumption of existing machine learning methods is that the data distribution of training data and unseen testing data is the same. When applying the machine learning model trained from cell line *in vitro* data to patient *in vivo* samples, the performance could significantly deteriorate due to the data distribution shift. Current efforts in solving the OOD problem include domain adaptation and meta-learning. Many domain adaptation methods have been proposed in computer vision and natural language processing. However, their application to aligning *in vitro* with *in vivo* data could be sub-optimal due to noisy and heterogeneous nature of omics data. An adversarial deconfounding autoencoder (ADAE) was proposed to facilitate the domain adaptation of gene expression profiles [4], but ADAE has not been tested for translating *in vitro* data to *in vivo* data. A meta-learning approach named TCRP has recently been proposed [5] to improve the transferability of predictive drug response models from *in vitro* screens to *in vivo* settings. However, TCRP still requires a certain number of patient data for each drug tested to train the predictive model. It is often infeasible to obtain such data, especially for new drugs. Thus the actual application of TCRP to clinic is limited. It can be only applied to the scenario of few-shot learning but not zero-shot learning. Most relevant to this work, Jia et al. has applied Variational Autoencoder (VAE) pre-training followed by Elastic Net supervised training (VAEN) to learn cell line models and applied them to impute *in vivo* drug response [6]. However, VAEN is not optimized to reliably transfer cell line data to patient samples and disentangle confounding factors [6] due to the fundamental limitation of VAE.

The unsolved question is how we can robustly predict individual patient response to a new drug that has never been tested in patients only using *in vitro* drug screens in the setting of zero-shot learning. To address this problem, we proposed a Context-aware Deconfounding Autoencoder (CODE-AE). In CODE-AE, we devised a self-supervised (pre)training scheme to construct a feature encoding module that can be easily tuned to adapt to the different down-stream tasks. We leverage both unlabeled cell line and tissue samples for the self-supervised (pre)training of the encoder. The unique features of CODE-AE are that it can extract both common biological signals shared by incoherent samples and private representations unique to them, and separate confounding factors from them. CODE-AE allowed us to generalize existing cell line omics data for the robust prediction of *in vivo* patient-specific response to new drugs in the setting of zero-shot learning, a critical component for patient-specific drug screening and personalized medicine.

To show the performance lift achieved by CODE-AE, we performed exhaustive comparative studies on CODE-AE (and variants) and other competing methods over the breast cancer patient-derived tumor xenograft *ex vivo* PDTC dataset [7]. Moreover, to demonstrate the potential of CODE-AE in personalized medicine, we apply CODE-AE to predicting chemotherapy resistance for patients in vivo, which is a significant obstacle to effective cancer therapy.

Lack of effective personalized chemotherapy tailored to individual patients often leads to unnecessary suffering and reduces the chances of patient’s overall survival. Our extensive studies show that CODE-AE effectively alleviates the out-of-distribution problem when transferring the cell line model to patient samples, significantly outperforms state-of-the-art methods AD-AE [4], TCRP [5], and VAEN [6] that are specifically designed for transcriptomics data as well as other popular domain adaptation methods Variational autoencoder (VAE) [8], Denoising autoencoder (DAE) [9], Deep Coral [10], and Domain Separation Network (DSN) [11] in terms of both accuracy and robustness. Using CODE-AE, we screened 50 drugs for 9,808 cancer patients. The *in vivo* drug screening not only further validated CODE-AE but also discovered novel personalized anti-cancer therapies and drug response biomarkers. Thus CODE-AE provides a useful framework to take advantage of rich *in vitro* omics data for developing generalized clinical predictive models.

## 2 Results and Discussion

### 2.1 Overview of CODE-AE

As illustrated in Figure 1, given the gene expression profiles of labeled cell lines and unlabeled patients as input, CODE-AE learns a nonlinear embedding function. The embedding function projects a high-dimensional expression profile of each cell line or patient to a low-dimensional vector, which distinguishes biological meaningful signals from confounding factors and transform the embeddings of cell lines and patients into the similar distribution. The embedding function is learned using both labeled and unlabeled data. Thus, CODE-AE is able to generalize to unlabeled data. Furthermore, aligning the distributions of cell line and patient embeddings across labeled and unlabeled samples can alleviate the OOD problem.

**Figure 1:**
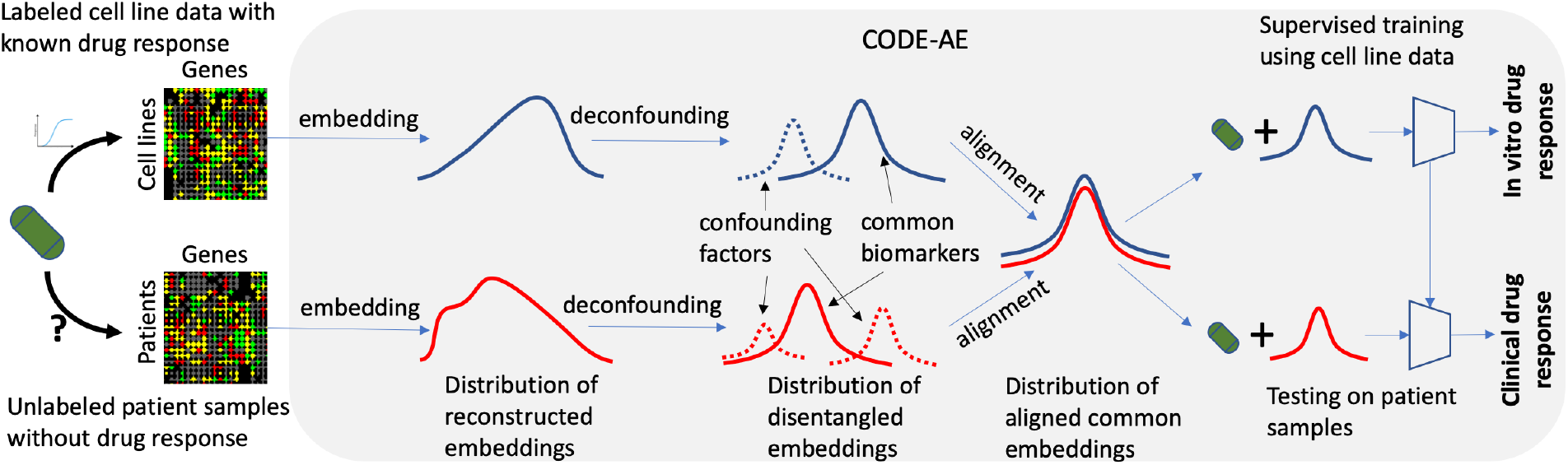
Illustration of CODE-AE method. Given labeled cell line drug response data, the aim of CODE-AE is to predict individual patient clinical response to drugs that has been tested in the cell line model but never been tested in the patient. Conceptually, CODE-AE consists of four steps. (1) The gene expression profile of both cell lines and patients are mapped into an embedding space. (2) Confounding factors are disentangled from intrinsic biomarkers in the embedding. (3) The distribution of embeddings of patients is aligned with that of cell lines. (4) A supervised model is trained based on the deconfounded and aligned embedding of cell lines, and tested using deconfounded and aligned embedding of patients.

Algorithmically, CODE-AE pretrains the neural network using an autoencoder that minimizes a data reconstruction error (See Methods). The pretraining step is useful for generalization to an unlabeled dataset. The brief architecture of CODE-AE is shown in Figure 2. Different from conventional autoencoders such as VAE, CODE-AE has two unique features. Firstly, it learns shared signals between the cell line data and the patient data as well as private signals that are unique to the cell line and the patient. The rationale is to disentangle common biological signals between data sets from context-specific patterns that overwhelm drug response biomarkers [5]. Secondly, CODE-AE regularizes the embeddings of cell lines and patients to have their distributions be similar. In this way, the knowledge learned from the cell line model can be transferred to patients. We test three regularization methods: simple concatenation of cell line and patient embeddings (CODE-AE-Base), minimization of their MMD loss (CODE-AE-MMD), and minimization of their adversarial loss (CODE-AE-ADV). After the unsupervised pretraining, a supervised drug response model can be trained from the aligned common embedding using the labeled cell line data. When a new patient comes, the drug response can be predicted from the trained cell line model based on the pretrained common embedding of the patient. Thus, CODE-AE does not need to use any labeled patient samples to construct the predictive model.

**Figure 2:**
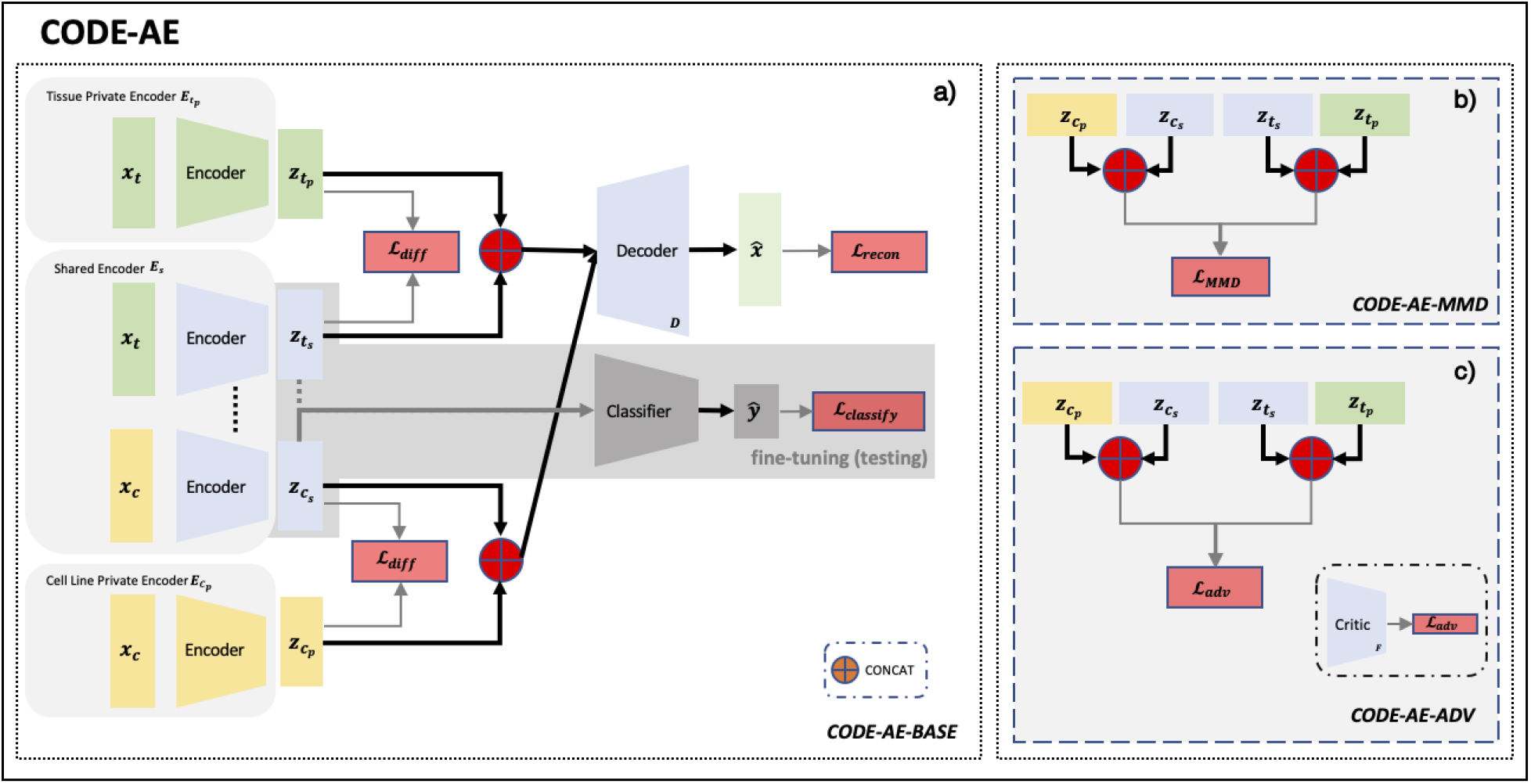
COntext-aware Deconfounding AutoEncoder (CODE-AE) framework. a) CODE-AE Base architecture: A layer-tying shared encoder **E**_*s*_ learns to map both cell line and tissue samples to deconfounded common intrinsic biological signals. Private encoders **E**._*p*_ learn to represent cell line/tissue context-specific information as private embeddings. A shared decoder **D** reconstructs the input samples through the concatenation of private and shared embeddings and the reconstruction quality is measured with 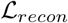. The private and shared embeddings are pushed apart through soft subspace orthogonality loss 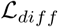. The shared encoder **E**_*s*_ appended with an additional classifier network will be trained during fine-tuning and perform inference during testing phase. b) CODE-AE-MMD: A variation of CODE-AE-BASE where the concatenation of private and shared embeddings are kept similar via optimizing 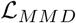. c) CODE-AE-ADV: A variation of CODE-AE-BASE where the concatenation of private and shared embeddings are kept similar via optimizing 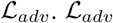 is in the form of min-max optimization between a critic network **F** and encoder components.

### 2.2 The optimal configuration of CODE-AE variants

We first studied the performance difference of CODE-AE variants in the combination of different configurations of the learning paradigm. In particular, the configuration choices we explicitly explored include (1) whether to include a hidden representation normalization layer, (2) use of only the shared representation or concatenation of private and shared representation for the downstream task, and (3) loss function for determining the similarity of tissue embeddings to cell line embeddings. We evaluated the performance of these CODE-AE variants using PDTC test dataset. As shown in Table 1 and Supplemental Figure S1, the overall best performing CODE-AE variant is the CODE-AE-ADV with hidden representation normalization, aligned cross-domain features over shared representation, and adversarial loss. Hidden representation normalization can avoid embeddings being pushed towards meaningless zero-valued vectors by soft orthogonality loss. The better performance achieved by using only the shared representation in downstream tasks is aligned with our assumption that shared representation is affluent with transferable deconfounded biological meaningful information. We will only compare CODE-AE-ADV with other baseline models and apply it to actual prediction tasks in the following sections.

**Table 1:**
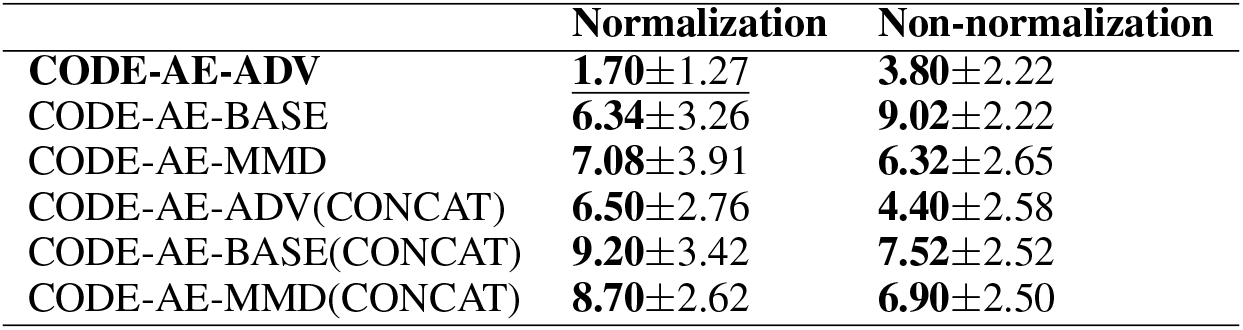
Average rank of CODE-AE variants on PDTC dataset

|  | Normalization | Non-normalization |
| --- | --- | --- |
| **CODE-AE-ADV** | **1.70±1.27** | **3.80±2.22** |
| CODE-AE-BASE | 6.34±3.26 | 9.02±2.22 |
| CODE-AE-MMD | 7.08±3.91 | 6.32±2.65 |
| CODE-AE-ADV(CONCAT) | 6.50±2.76 | 4.40±2.58 |
| CODE-AE-BASE(CONCAT) | 9.20±3.42 | 7.52±2.52 |
| CODE-AE-MMD(CONCAT) | 8.70±2.62 | 6.90±2.50 |

### 2.3 CODE-AE-ADV alleviates the out-of-distribution problem on transferring cell line models to patient data

We used the shared encoder from pre-trained CODE-AE-ADV to generate the new representations for *in vivo* TCGA patient samples and in vitro CCLE cell line samples. To inspect how well the embeddings of cell line data and patient samples are aligned, we generated tSNE plots to visualize their embeddings, as shown in Figure 3. The embeddings of TCGA and CCLE samples largely overlap in tSNE manifolds. It indicates that CODE-AE-ADV is effective in aligning cell lines and patients’ representations. As a comparison, the low-dimensional representations of CCLE and TCGA data are clearly separated when using original gene expression profiles or vanilla autoencoder. Thus, CODE-AE-ADV is more effective in addressing OOD problem than the embedding algorithms that are used by the state-of-the-art method VAEN [6].

**Figure 3:**
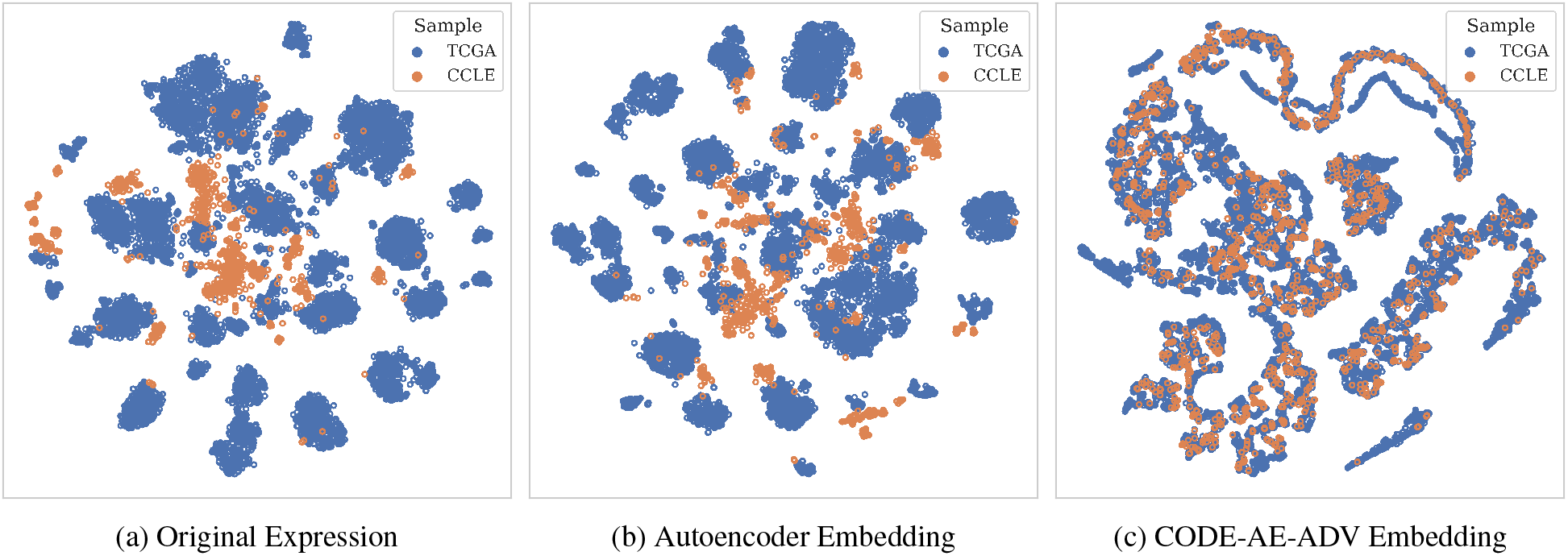
tSNE plots of a) original expression b) embeddings generated by standard autoencoder c) embeddings generated by CODE-AE-ADV

### 2.4 CODE-AE-ADV significantly outperforms baseline models when predicting *ex vivo* drug responses

We then compared CODE-AE-ADV with baseline models using *ex vivo* drug response data from PDTC. As shown in Table 2, CODE-AE-ADV is overall the best performer for PDTC test dataset. The second best performer is ADAE that is specially design to remove confounders [4]. Interestingly, the state-of-the-art domain adaptation methods DSN, CORAL, and DANN in computer vision and natural language processing do not perform well, even worse than the standard VAE. This observation suggests that omics data could be fundamentally different from images and human languages. Specially designed deep learning model is needed to address the challenges in omics data integration and predictive modeling. Among these domain adaptation methods, DSN performs the best, suggesting the importance in disentangling shared and private information between cell line and patient samples. It is not surprising that models incorporated unlabeled pre-training clearly outperform the ones without. TCRP on average is the best performer among models without unlabeled pre-training.

**Table 2:**
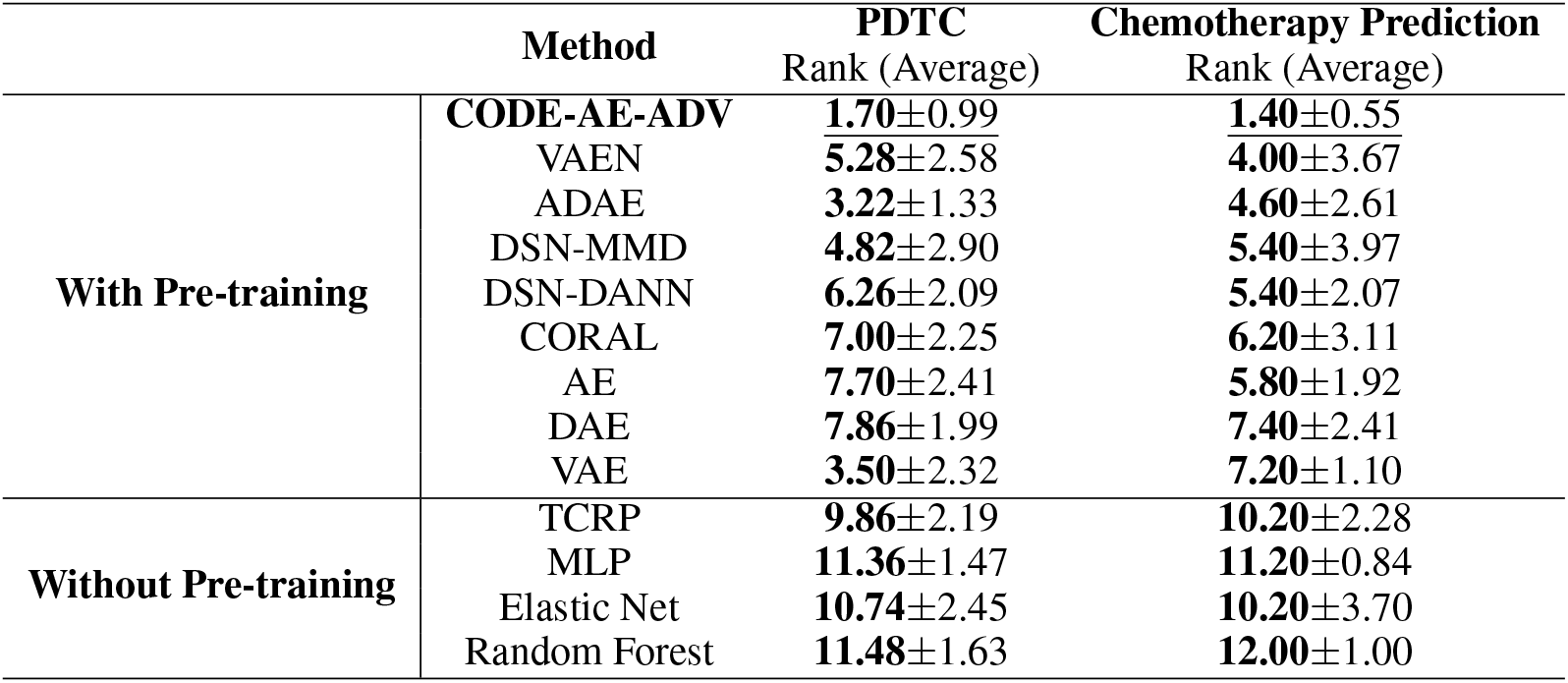
Average rank of different methods on PDTC test dataset and patient chemotherapy response prediction

Figure 4 shows the drug-wise performance of each algorithm. Among 50 drugs tested, CODE-AE-ADV ranks as the best, the second best, and the third best for 29, 11, 7 times, respectively. It performs the best for three drugs BX795, Obatoclax Mesylate, and Axitinib with the AUROC above 0.8. The worst ranked drugs by CODE-AE-ADV are PD0325901, SB216763 and AZD7762. The reason for this performance disparity is unclear and worth further investigation.

**Figure 4:**
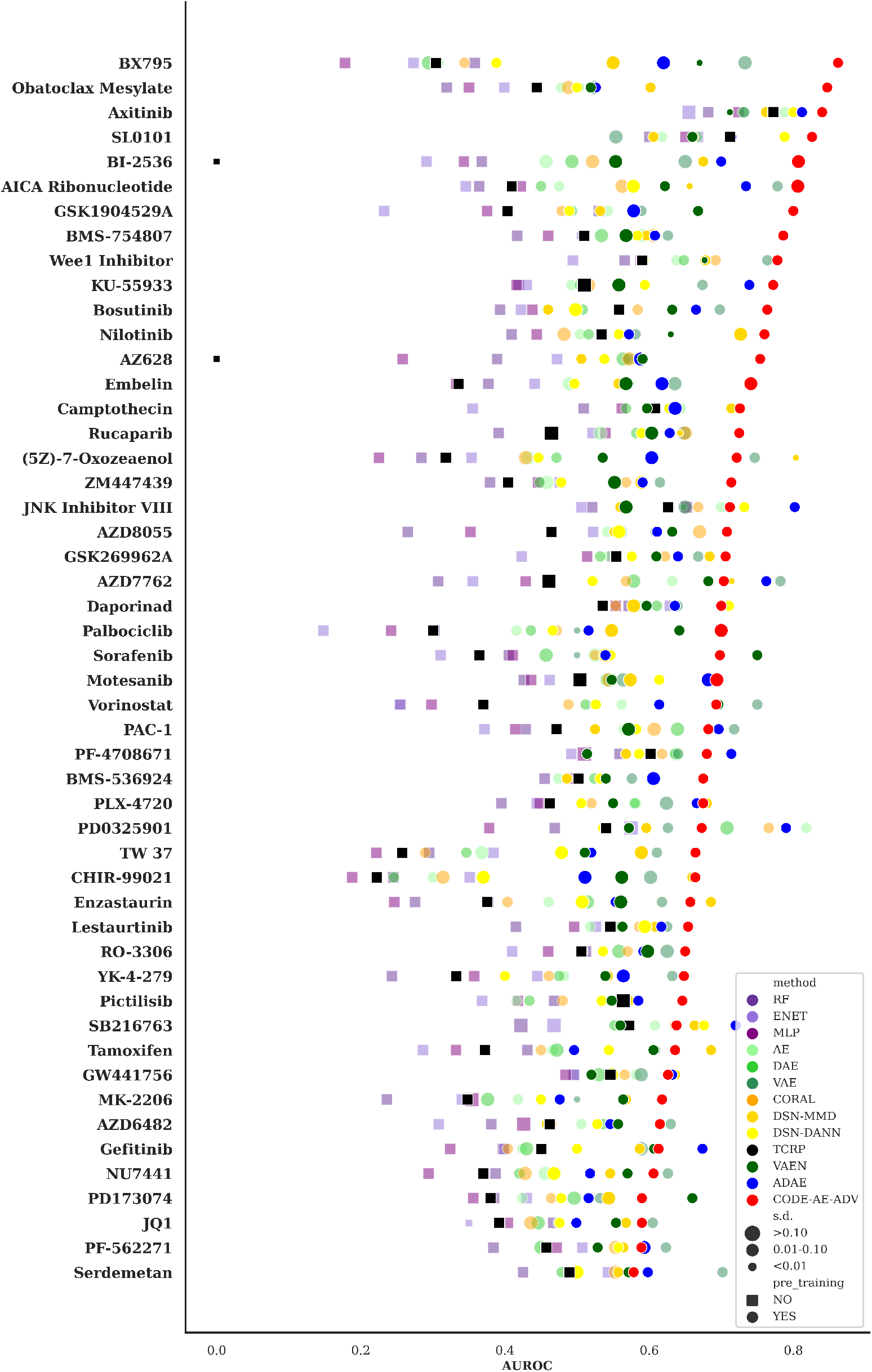
Performance comparison of PDTC drug response classification

### 2.5 CODE-AE-ADV significantly outperforms baseline models when predicting patient chemotherapy resistance

We further evaluated the performance of CODE-AE-ADV for predicting clinical chemotherapy resistance in two aspects: either a lack of reduction in tumor size following chemotherapy or the occurrence of clinical relapse after an initial “positive response to treatment” [12] as detailed in Method section. The results are shown in Figure 5. Consistent with the results from the PDTC data set, CODE-AE-ADV significantly outperforms baseline models. CODE-AE-ADV achieved the highest value of AUROC in 5 out of 7 cases with statistically significant (p-value ≤ 0.05) performance gain, and second highest in the other 2 but without statistically significant difference from the best one. CODE-AE-ADV is overall the best performer in this task as shown in Table 2. VAEN performs relatively well in the case of relapse days after treatment, but much worse than CODE-AE-ADV, ADAE, and TCRP when evaluated by the clinical diagnosis. Furthermore, CODE-AE-ADV significantly outperforms ADAE by a large margin in all cases. This observation further supports that CODE-AE-ADV can enhance the signal-to-noise ratio in the biomarker identification because the major difference between CODE-AE-ADV and ADAE is to disentangle shared and private embeddings between cell lines and patient tissues. TCRP is inferior to the other methods that take advantage of the pre-training using unlabeled data, demonstrating the importance of the pre-training in few-shot and zero-shot learning.

**Figure 5:**
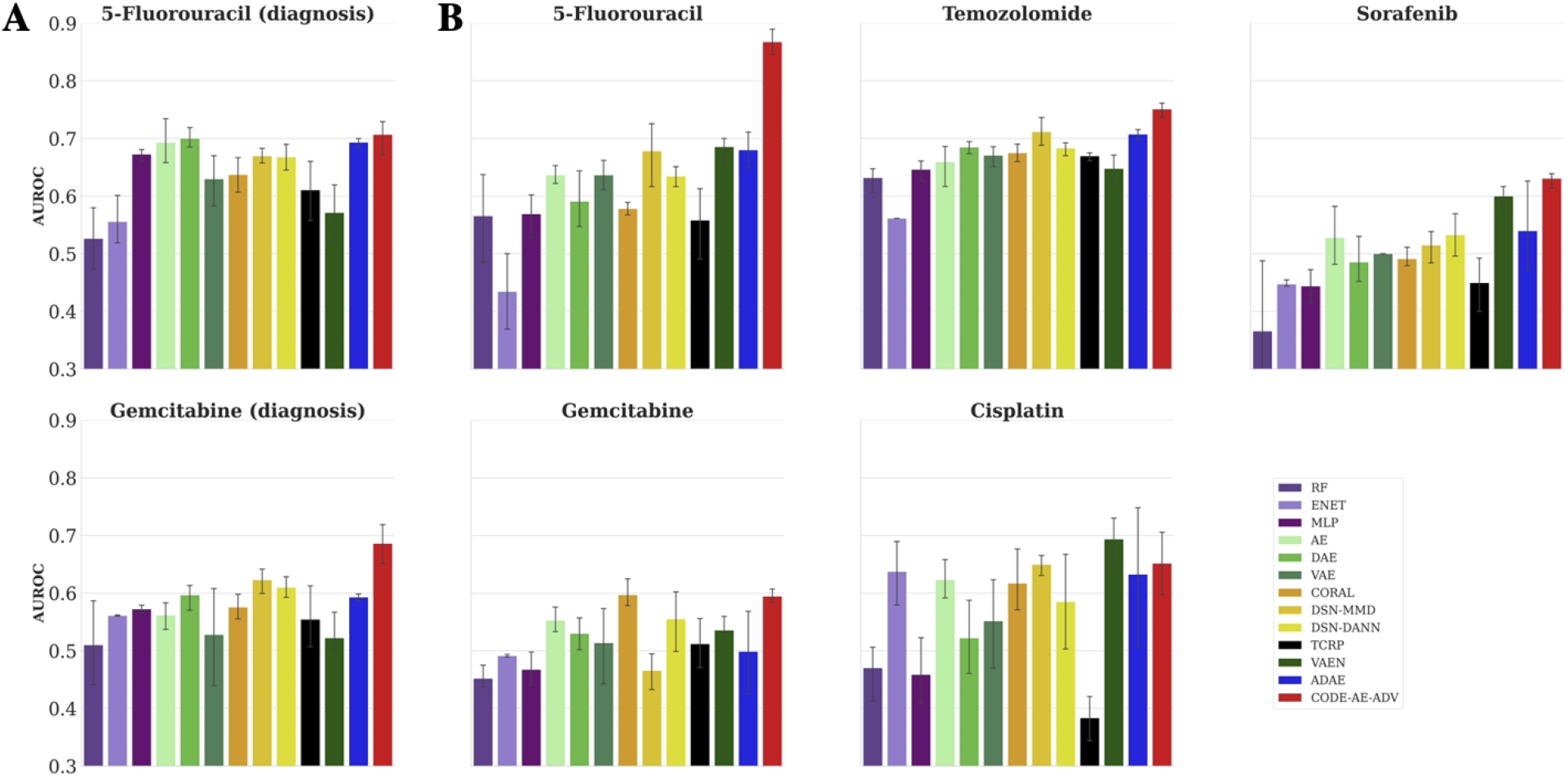
Performance comparison of patient chemotherapy response prediction based on (A) clinical diagnosis, and (B) relapse days after treatment. When classifying the clinical diagnosis for drug 5-Fluorouracil and Gemcitabine, CODE-AE-ADV outperforms the second best performer statistically significant with a p-value of 0.0399 and 0.0290, respectively. For the classification based on relapsed days after first treatment, CODE-AE-ADV significantly outperforms the second best performer for drug 5-Fluorouracil, Temozolomide, Sorafenib with a p-value of 0.0028, 0.0483 and 0.0420, respectively. In the case of drug Gemcitabine and Cisplatin, CODE-AE-ADV ranked the second best, while its performance difference with the respective best performing methods is not statistically significant with a p-values of 0.8746 and 0.3102, respectively.

### 2.6 CODE-AE-ADV is successful in deconfounding omics data

To show that CODE-AE-ADV can generate transferable embedding through deconfounding uninteresting confounders while preserving true biological signals present in expression data even outside the in-vivo and in-vitro setting. We selected the gene expression data sets used in ADAE [4] to perform a similar evaluation process. Specifically, we chose the brain cancer expression data set with gender information as confounding factors and brain cancer subtype classification as target downstream tasks. We first performed encoder training with all unlabeled gene expression profiles regardless of gender. For ADAE [4] and CODE-AE, we selected the binary gender variable as the deconfounding target. After encoder training, we generated the latent embedding for all original gene expression profiles using different encoders. Then, we built elastic net classifiers for cancer subtype prediction using the latent embedding of samples of one gender to predict the other gender samples. Following the evaluation procedure described in [4], the classification performance measured in the area under the precision-recall curve (AUPRC) as well as area under the receiver operating curve (AUROC) of ten-fold cross-validation was reported in Table 3. Besides, we performed a two-sample t-test on the average performance between CODE-AE-ADV and the best non-CODE-AE method in each setting, and its results are shown on the last row of Table 3. We observed the same trends as those in the drug resistance prediction. Using the model built from female data to predict male data, CODE-AE-ADV significantly outperforms ADAE, the second-best performer measured by both AUROC and AUPRC. When applying the model trained from male data to predict female data, the performance of CODE-AE-ADV is slightly worse than CORAL, but the difference is not statistically significant. Both CODE-AE-ADV and CORAL significantly outperform the state-of-the-art deconfounding method ADAE (p-value ≤ 0.05). Additionally, two other observations from the drug response experiments hold. Disentangling common and private features of different data modalities is essential for cell line to tissue transfer learning, and adversarial loss is more effective than MMD loss.

**Table 3:**
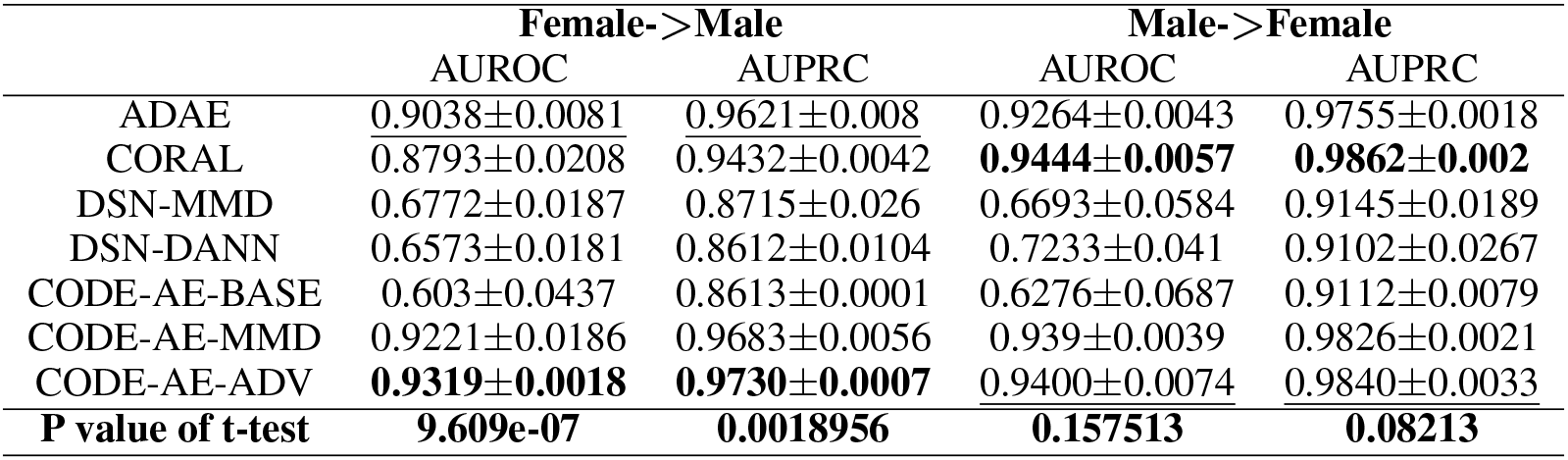
Performance comparison on cancer subtype prediction with the gender as a confounding factor. The best and the second best performances are highlighted and underlined, respectively.

|  | Female->Male |  | Male->Female |  |
| --- | --- | --- | --- | --- |
|  | AUROC | AUPRC | AUROC | AUPRC |
| ADAE | 0.9038±0.0081 | 0.9621±0.008 | 0.9264±0.0043 | 0.9755±0.0018 |
| CORAL | 0.8793±0.0208 | 0.9432±0.0042 | **0.9444±0.0057** | **0.9862±0.002** |
| DSN-MMD | 0.6772±0.0187 | 0.8715±0.026 | 0.6693±0.0584 | 0.9145±0.0189 |
| DSN-DANN | 0.6573±0.0181 | 0.8612±0.0104 | 0.7233±0.041 | 0.9102±0.0267 |
| CODE-AE-BASE | 0.603±0.0437 | 0.8613±0.0001 | 0.6276±0.0687 | 0.9112±0.0079 |
| CODE-AE-MMD | 0.9221±0.0186 | 0.9683±0.0056 | 0.939±0.0039 | 0.9826±0.0021 |
| CODE-AE-ADV | **0.9319±0.0018** | **0.9730±0.0007** | 0.9400±0.0074 | 0.9840±0.0033 |
| P value of t-test | **9.609e-07** | **0.0018956** | **0.157513** | **0.08213** |

### 2.7 Application of CODE-AE-ADV to personalized medicine

To further validate CODE-AE-ADV with patient data and demonstrate its utility in personalized medicine, we applied CODE-AE-ADV (per-drug) trained with CCLE data to screen 50 drugs for 9,808 cancer patients from TCGA. Our major findings are summarized below.

#### 2.7.1 Gene expression differential analysis of drug target verifies predicted patient drug responses from CODE-AE-ADV

We first verify our predictions by checking the association of our predicted drug response with the gene expression values of drug targets. If the predicted patient response on the targeted therapy is correlated with the drug target, it provides the validation of our prediction. We select the top 5% predicted drug sensitive patients as our responsive patient set and bottom 5% as the resistant patient set. We found that 47 out of 50 targeted therapies are statistically significantly (p-value ≤ 0.05) associated with the drug target expression (Supplemental Table S1 and Supplemental Figure S2). For instance, three drugs targeting RTK signaling pathway, Sorafenib, AMG-706 and Axitinib, consistently have strong association between predicted drug response and their target gene expression (Supplemental Figure S3). The targets of these drugs include KIT, KDR, PDGFRA and PDGFRB, which are all key components in the RTK signaling pathway. The same phenomenon could also be noticed for other targeted drugs. This indicates that CODE-AE-ADV could well capture the drug mode of action.

#### 2.7.2 Gene Set Enrichment Analysis of predicted patient drug response gene expression profile identifies drug response biomarkers

We further conducted a Gene Set Enrichment Analysis (GSEA) for the gene expression profile of sensitive and insensitive patients for each drug to elucidate biological mechanisms underlying the predicted drug-patient association. For example, three enriched gene sets were identified for the Gefitinib-sensitive patients including down-regulated genes when KRAS is overexpressed. On the other hand, the up-regulated genes for tissues with overexpressed mutated KRAS are enriched for the Gefitinib-resistant patients. It has been well known that patients with mutated KRAS have significantly worse prognosis to EGFR-inhibition cancer therapy [13, 24, 14]. Our results are consistent with these observations. For IGF-IR inhibitor GSK1904529, the top-rank gene sets in the resistant patients include up-regulated genes by the over-expression of AKT, MET or ERBB genes. Because receptor tyrosine kinases, like IGF-IR, MET and ERBB, can regulate PI3K/AKT/mTOR pathway, JAK/STAT pathway and Ras/Raf/Mek/Erk pathway[15], whose functions overlap with each other. Therefore, the effect caused by inhibition of IGF-IR could be compensated by other kinases that are up-regulated in GSK1904529-resistant patient tissues. Another statistically significant example is TAK1 inhibitor (5Z)-7-Oxozeaenol. We found that the down-regulated and up-regulated gene set in mutated P53 cell line were enriched in the (5Z)-7-Oxozeaenol-sensitive and -resistant patients, respectively. This indicates that P53 might play an important role for patient response to (5Z)-7-Oxozeaenol. Interestingly, it is reported that TAK1 signaling pathway can regulate p53 [22]. Thus, p53 expression level could be a drug response biomarker for (5Z)-7-Oxozeaenol. The predicted gene set enrichment of several other drugs (AXD6482, SL0101-1, AICAR, and NU7441) is also supported by existing experimental evidences.

**Table 4:**
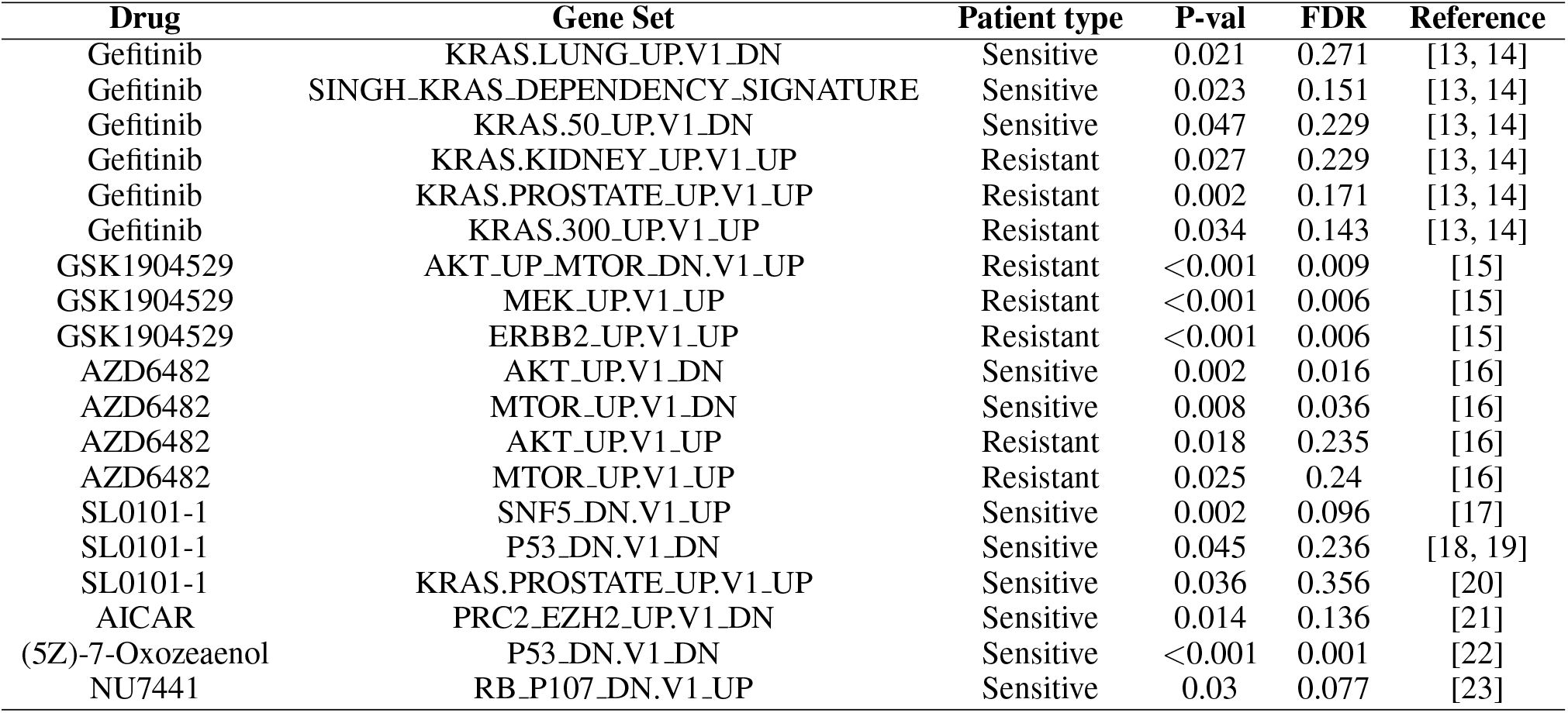
Summarized GSEA results of predicted patient drug response gene expression profile

#### 2.7.3 Clustering analysis of drug response profiles reveals novel connections between drugs and between tumor types

We performed clustering analysis on the predicted drug response matrix as shown in Figure 6, within which the rows and columns are corresponding to drugs and patients, respectively. Specifically, we grouped 50 drugs into 16 clusters with Spectral Co-clustering [25] and stratified 9,808 patients into 33 clusters with Kmeans++ clustering [26]. Notably, each patient cluster is dominated by one of the patient tumor types, suggesting the clustering is clinically meaningful. However, patients with different tumor types could be grouped into the same cluster. For example, the cluster shown in Figure 6 consists of adrenal gland tumors besides the brain tumor. It implies that clinical diagnosis itself may not identify the best therapy for the patient. The stratification of patients based on the drug response profile could be a more effective strategy for personalized medicine. Detailed inspection of drug clusters reveal that drugs with different primary targets may lead to the similar drug response. For example, the largest drug cluster consists of 15 drugs (SB 216763, AZD8055, BMS-754807, PF-4708671, JNK Inhibitor VIII, PLX4720, AG-014699, AMG-706, TW 37, PD-0325901, GDC0941, GW 441756, Bosutinib, PAC-1 and BI-2536). These drugs are designed to target several pathways, notably, RTK signaling, IGF1R/PI3K signaling, and ERK/MAPK signaling. On the one hand, these pathways are strongly inter-connected. RTK activates PI3K or MAPK pathways [27]. On the other hand, it is likely that multi-target bindings may contribute to the similar clinical response of these drugs [28].

**Figure 6:**
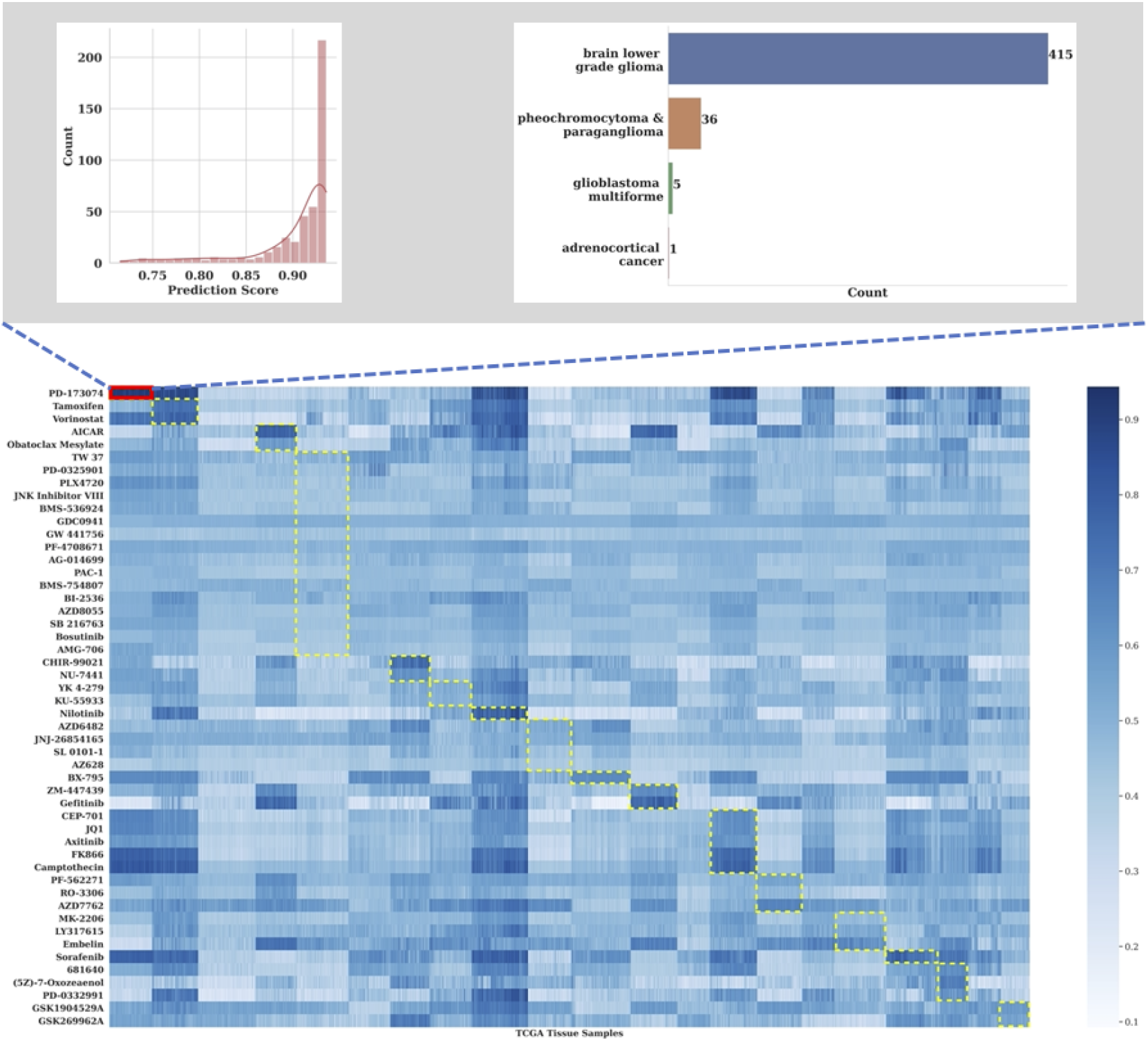
Co-clustering results on complete patient drug response prediction. Yellow dashed boxes indicate the identified clusters. Top two figures demonstrate the composition of cluster 0 which include only drug PD173074. Top left figure shows the prediction score distribution within this cluster, which indicates the high sensitivity of patients within this cluster under the treatment of PD173074. Top right figure shows the tissue origin of samples within this cluster. 415/457 samples are from brain lower grade glioma tissue.

#### 2.7.4 CODE-AE-ADV identifies precision anti-cancer therapy

We ranked drugs based on their mean predicted responses for each patient cluster (Supplemental Table S2) to identify the precision anti-cancer therapy. Intriguingly, we found that the glioblastoma (GBM) patient shows high sensitivity to PD173074, a fibroblast growth factor receptors (FGFR) inhibitor. Several studies have shown that even though FGFR gene mutation is rare in GBM, the FGFR is crucial in the regulation of many downstream functions, including cell survival, cell proliferation and cytoskeletal regulation etc [29]. The FGFR3-TACC complex has also been demonstrated to be carcinogenic in GBM [30]. Given that the FGFR inhibitor has been determined to hinder the pediatric glioma cells growth, PD173074 could be a potential anti-cancer therapy for GBM [31]. Liver cancer and acute myeloid leukemia (AML) are also predicted to be sensitive to PD173074. Experimental studies have shown that FGFR4 is over-expressed in livers and indicated that its up-regulation contributes to the progression of hepatocellular carcinoma [32]. For AML, previous study has demonstrated the FGFR inhibitors could suppress the leukemogenesis, especially for FGFR1 overexpressed AML [33]. Thus, our top ranked predictions are supported by existing experimental and clinical evidences. CODE-AE-ADV also predicted that pheochromocytoma & paraganglioma and ovarian serous cystadenocarcinoma are sensitive to PD173074. In addition to PD173074, several other drugs, notably, Camptothecin, Vorinostat, and Sorfenib, were predicted to be effective on multiple tumors. These predictions warranty further experimental validations.

## 3 Conclusion

In this paper, we introduce a new transfer learning framework CODE-AE to predict individual patient drug response from a supervised neural network model trained from cell line data. Extensive benchmark studies demonstrate the advantage of CODE-AE over the state-of-the-art in terms of both accuracy and robustness. The performance gain of CODE-AE mainly comes from (1) the unsupervised learning that combines unlabeled data from both cell lines and patient samples, (2) separation of shared common features cross cell lines and patient samples with unique embedding for cell lines or patients, and (3) adversarial training to optimize the similarity and difference between incoherent data sets. CODE-AE could be further improved in several directions. In contrast with cell line data from a pure population of cells, patient tissue data are mixtures of normal, abnormal, and infiltrated immune cells. We can further improve the CODE-AE by the deconvolution of patient gene expression data. We only use transcriptomics profiles to build the predictive model in this study. We can integrate additional omics data such as somatic mutations and copy number variants in the framework of cross-level information transmission [34]. Finally, we only apply CODE-AE to cancers. It will be interesting to test the performance of CODE-AE in other diseases besides cancers, which even do not have a large number of cell line data. In principle, CODE-AE can be applied to other transfer learning tasks with two data modalities with shared and unique features.

## 4 Methods

### 4.1 Our Approach: COntext-aware DE-confounding AutoEncoder (CODE-AE)

We proposed a novel CODE-AE to generate biologically informative gene expression embeddings to transfer knowledge from in-vitro data into patient samples. CODE-AE employed the standard auto-encoder as the backbone to leverage the unlabeled gene expression data sets. Inspired by the work on factorized latent space [35] and domain separation network [11], we encoded the samples (from cell lines or tumor tissues) into two orthogonal embeddings, namely private embeddings and shared embeddings. The first one is designed to separate the context-specific signals that overwhelm the common biomarkers. The latter contains the deconfounded common intrinsic biological signals used to transfer knowledge across cell lines and tissues.

#### 4.1.1 CODE-AE Base

As shown in Figure 2, the C DE-AE takes ex ress n vectors from *in vitro* cell lines and patient tumor tissue samples as input. Let 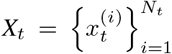 and 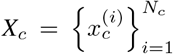 represent the unlabeled data set of *N*_*t*_ patient tumor tissue samples and *N*_*c*_ *in vitro* cancer cell line samples, respectively. Each sample *x* will be encoded into two separate embeddings through its corresponding cell line or tissue private encoder **E**._*p*_ and also the weight-sharing encoder **E**_*s*_. The concatenation of these two embeddings of each sample is expected to be able to reconstruct the original gene expression vector *x* through a shared decoder **D**, and the reconstruction is done as,

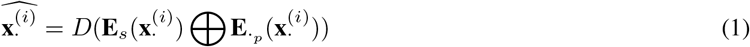

where **x**.^(*i*)^ represents the input gene expression profile, 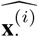 is the corresponding reconstructed input sample through the autoencoder component. ⊕ stands for the vector concatenation operation. We measure the quality of autoencoder reconstruction through the mean squared error between the original samples and the reconstruction output as below,

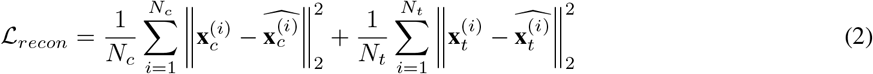

In our formulation, we factorized each sample’s latent space into two different subspaces to capture both domain specific and common information separately. To minimize the redundancy between the factorized latent spaces, we included an additional penalty term, 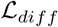 in the form of orthogonality constraint. The difference loss 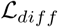 is applied to both cell line and tissue samples and encourages the shared and private encoder to encode different aspects of the inputs. We define the loss via soft subspace orthogonality constraint as below,

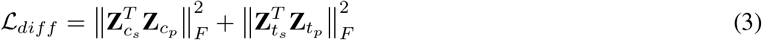

where **Z**._*s*_ are the embedding matrices whose rows are the shared embedding for cell line or tissue samples, while **Z**._*p*_ are the embedding matrices whose rows are the private embedding for cell line or tissue samples. It is obvious that 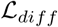 tends to push the embeddings to meaningless all-zero-valued vectors. To avoid such scenario, we append an additional instance normalization layer after the output layer of each encoder to avoid embeddings with minimal norm. Lastly, the loss for CODE-AE-BASE is defined with the weighted combination between 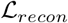 and 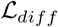 as below,

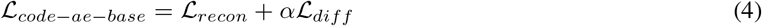

where *α* is the embedding difference loss coefficient.

#### 4.1.2 CODE-AE Variants

With CODE-AE-BASE, we could split cell line or tissue sample’s inherent information into the private and shared streams. However, in our baseline experiments, we often found that it was sub-optimal or demonstrated varied performance. Thus, we proposed two variants that showed better and generally more stable performance. Under the CODE-AE framework, for each input sample, CODE-AE factorized it into two orthogonal embeddings. The concatenation of these two embeddings is considered as the new representation of the original input. Given that all samples in our consideration are gene expression profiles regardless of cell line or patient, we assumed that the new representation of original input in the factorized latent space close to each in terms of distributional differences. Hence, we incorporated additional feature alignment component into the CODE-AE-BASE framework. Specifically, the distributional difference of the concatenated representation of private and shared embeddings from both cell line and tumor tissue samples are minimized via the following two approaches.

##### CODE-AE-MMD

The first variant, named CODE-AE-MMD, utilized the well known maximum mean discrepancy [36] as the distance measurement between the latent representation of cell line and tissue samples. Maximum Mean Discrepancy (MMD) loss [36] is a kernel-based distance function between samples from two distributions. In particular, we used an approximate version of exact MMD loss in CODE-AE-MMD as below,

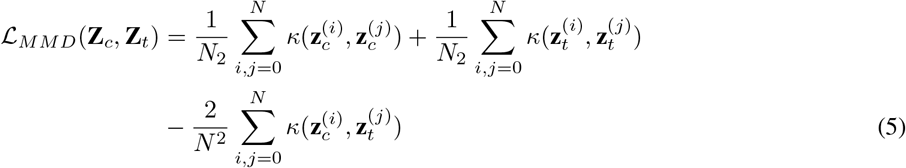

where **Z**_*c*_, **Z**_*t*_ are embedding matrices for cell line and tissue samples respectively, whose rows are the concatenations of each sample’s private and shared embedding. **z**.^(*i*)^, **z**.^(*j*)^ are the *i*-th or *j*-th samples’ corresponding embedding vectors. In practice, *N* will be the batch size. Accordingly, the loss of CODE-AE-MMD is given as below,

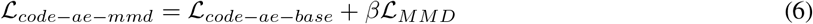

where *β* is the MMD loss coefficient.

##### CODE-AE-ADV

The second variant, CODE-AE-ADV, employed adversarial training to push the representations of cell line and tissue samples to be similar to each other. Specifically, w e a ppended a c ritic n etwork *F* t hat scores representations with the objective that consistently gives higher scores for representations of cancer cell line samples. The encoders for tissue samples are given an additional objective to generate the embedding that could fool the critic network to produce high scores. In this manner, critic network and tissue sample encoders will play a min-max game in the form of an alternative training schedule, which is adopted by Wasserstein generative adversarial networks [37]. To avoid unstable training commonly existing in alternative training schedules, instead of standard WGAN [37], we used the WGAN with gradient penalty [38]. Its affiliated loss terms are defined as below,

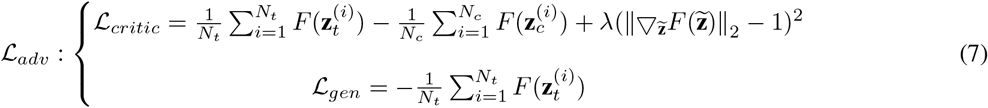

where **z** = **z**._*s*_ ⊕ **z**._*p*_ stands for new representation of input and 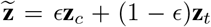 and *ɛ* ∼ **U**(0,1). A detailed CODE-AE-ADV learning procedure can be found in (Procedure 1).

**Procedure 1.**
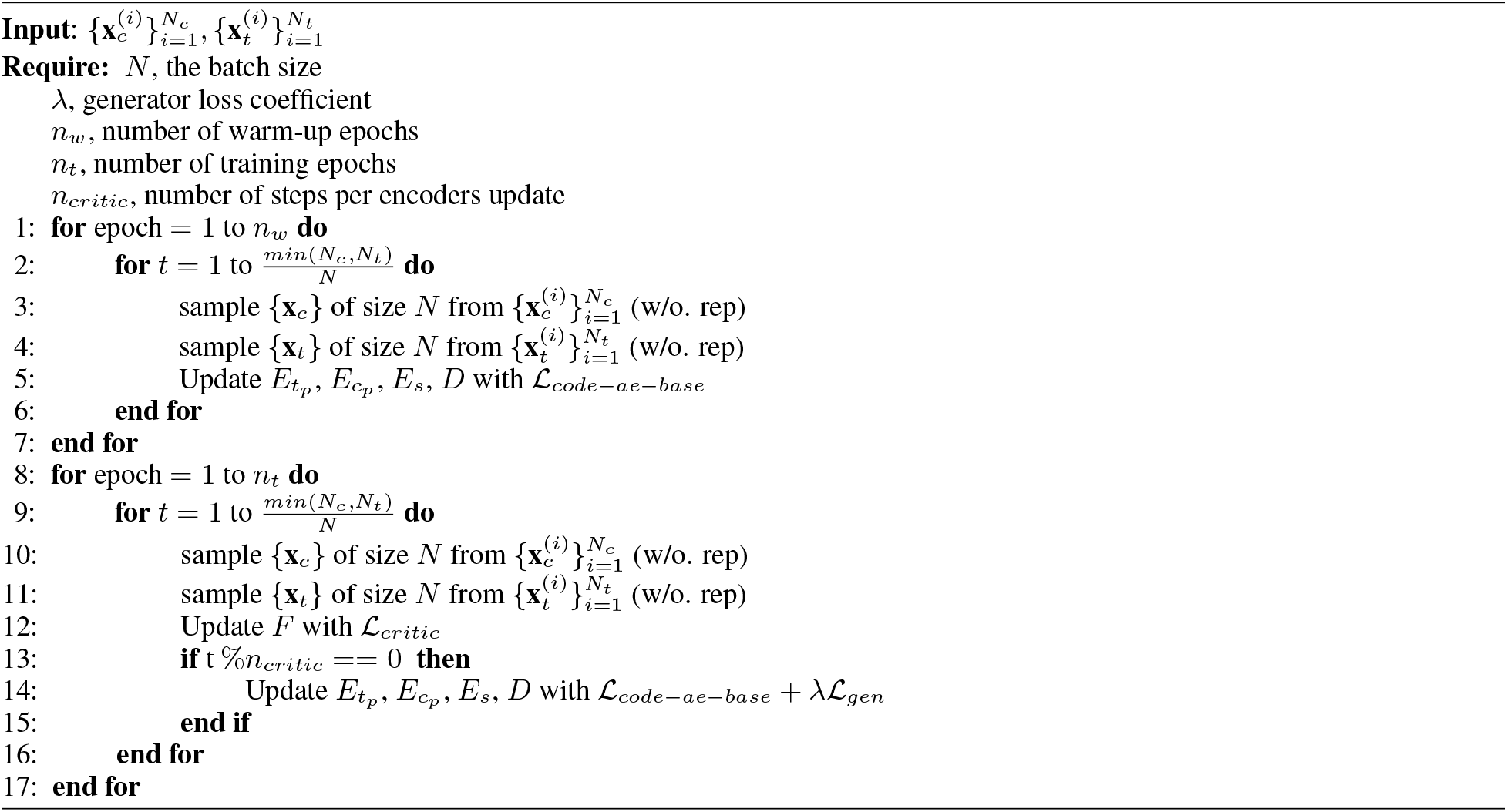
CODE-AE-ADV training

After the encoder training with unlabeled data as mentioned above, the shared encoder **E**_*s*_ could be used to directly generate the deconfounded biological meaningful embedding vectors or append a neural network module for specific downstream tasks. In the latter case, strategies such as gradual unfreezing and decayed learning rate schedule could be adopted to improve task-specific performance further, as shown in our following experiments.

### 4.2 Gene set enrichment analysis

For each drug in consideration, we select the top 5% predicted drug sensitive patients as our responsive patient set and bottom 5% as the resistant patient set. Then, we input each responsive/resistant patient tumor tissue gene expression profiles (20000 genes) and their respective predicted response label into GSEA[39, 40]. Our enrichment analysis is focused on the oncogenic signature gene sets. Each gene set includes a list of genes that are regulated after perturbation of some cancer related genes. For each drug, the statistically enriched gene set for sensitive and resistant patient tissues are used to be explored further.

### 4.3 Clustering analysis

We separated 50 drugs into 16 clusters with Spectral Co-clustering methods based on the predicted drug responses profiles. Then we applied Kmeans++ clustering to group 9,808 patients into 33 clusters. For each cluster, we averaged tissues drug response score profiles and then ranked the drugs based on the average scores. The higher score indicates that this cluster of patient is more sensitive to the drug.

### 4.4 Experiments Setup

#### 4.4.1 Data sets

##### Training Dataset

###### Unlabeled pre-training (*in vitro* and *in vivo*)

The unlabeled datasets used for encoder pre-training include cancer cell line and patient tumor tissue gene expression profiles. Specifically, we collected 1,305 cancer cell line samples with corresponding gene expression profiles from the DepMap portal [41] and 9,808 patient tumor tissue samples from the Xena portal [42]. All gene expression data are metricized by the standard transcripts per million base for each gene, with additional log transformation. In addition, we selected the top 1000 varied genes measured by the percentage of unique values in gene expression samples for cancer cell lines and tumor tissue samples separately. Then we combined the two sets of top 1000 varied genes as the input features. There are a total of 1426 genes in the feature set.

###### Labeled fine-tuning(*in vitro*)

The labeled dataset used for fine-tuning phase is collected from GDSC [43, 44]. GDSC recorded the cellular growth responses of cancer cell lines against a panel of drugs as the area under the drug response curve (AUC), which is defined as the fraction of the total area under the drug response curve between the highest and lowest screening concentration in GDSC. For each drug of interest, we first identified all cell lines with corresponding drug sensitivity measured in the area under the drug response curve (AUC) and then split these cancer cell lines’ sensitivity against this drug into binary labels, namely responsive or non-responsive (resistant). The categorization threshold is selected as the average AUC value of all available cell line drug sensitivity for each drug of interest.

##### Test Dataset

We evaluated the performance of CODE-AE in the setting of zero-shot learning, i.e., the unseen OOD data have never been used in training. It is a more difficult but more realistic scenario than the state-of-the-art method TCRP [5] in which a small set of OOD data was used during the training. Specifically, the predictive model for each drug of interest was learned only with the aforementioned *in vitro* dataset. While in testing time, we evaluated the model performance with the following *ex vivo* and *in vivo* labeled datasets that were not used in the training phase on the prediction task of drug response classification in pre-clinical and clinical scenarios, respectively.

###### Pre-clinical (*ex vivo*)

We used data from breast cancer PDTC [7] to evaluate the performance of drug response classification in a pre-clinical context. The previous study collected 83 human breast tumor biopsies and established human cell culture from these tumors with mice as intermediaries. Each of these human cell cultures was exposed to a list of drugs. From the list of drugs available in PDTC, we further selected 50 drugs with known protein targets for which cell-line responses had also been recorded in GDSC as drugs of interest. The drug sensitivity classification of each drug was considered as a separate learning task. Similar to the labeled GDSC dataset used during training, the PDTC responses were categorized into binary labels using PDTC AUCs, where the classification threshold is specified as the median AUC value of all available PDTC AUCs of each drug of interest.

###### Clinical (*in vivo*)

To evaluate the performance of drug response classification in a clinical context, we primarily consider a practical problem: predict chemotherapy resistance given gene expression profiles of patients while training the predictive model *only* using the gene expression profile of cancer cell lines.

Clinical chemotherapy resistance can be defined as either a lack of reduction in tumor size following chemotherapy or the occurrence of clinical relapse after an initial “positive response to treatment” [12]. Hence, we extracted data sets to assess these two aspects. The patient clinical drug response was acquired from a recent work [45], where patients’ clinical response records of two chemotherapy agents Gemcitabine and Fluorouracil from The Cancer Genome Atlas (TCGA) [46] were extracted. The patients were split into two groups: responders who had a partial or complete response and non-responders who had progressive clinical disease or stable disease diagnosis. Only patients on single-drug therapy through the entire duration of treatment were retained in the study.

In addition to using clinical diagnosis to indicate patients’ drug responses towards a particular drug, we extracted patients’ “new tumor events days after treatment” from TCGA [46] as the standard to divide patients into responders and non-responders. The median number of days of new tumor events was used as the threshold. Similar to the above data set from [45], we only included patients on single-drug therapy through the entire treatment duration in this test data set. For the list of drugs included in this test dataset, the drugs with more than 20 labeled samples are kept.

#### 4.4.2 Baseline models

We compared CODE-AE with the following base-line models that include unlabeled pre-training: VAEN [6], standard autoencoder (AE) [47], denoising autoencoder (DAE) [9], and variational autoencoder (VAE) [8] as well as representative domain adaptation methods including deep coral (CORAL) [10] and domain separation network (DSN) [11] of both MMD (DSN-MMD) and adversarial (DSN-DANN) training variants. Furthermore, we included a more recent adversarial deconfounding autoencoder (ADAE) [4] given its similar formation as DANN [48] and state-of-the-art performance in transcriptomics data sets. In addition, for CODE-AE variants, we also explored different configurations, such as with/without hidden layer normalization, performing a downstream task with concatenated representation, or shared representation in an ablation study with PDTC test dataset.

For fair comparisons, all the encoder and decoder trained in the experiments share the same architecture. Specifically, the hidden representation is of dimension 128. The encoders and decoder are 2-layer neural network modules of dimension (512, 256) and (256, 512), respectively, with the rectified linear activation function. Appended modules such as critic network in CODE-AE-ADV and classifier network used for fine-tuning are 2-layer neural networks of dimension (64, 32) with rectified linear activation, have one output node with linear activation in critic network, and sigmoid activation in classifier networks. Further, the loss weight terms in CODE-AE-MMD and CODE-AE-ADV are all specified as 1.0.

Moreover, for models that do not include unlabeled pre-training, we compared CODE-AE with TCRP[5] as well as vanilla neural network (denoted as MLP), elastic net classifier (denoted as EN), and random forest classifier (denoted as RF). TCRP incorporates model agnostic meta-learning technique and is the most successful method for predicting individual patient drug response from the cell line data so far.

#### 4.4.3 Training procedure

For models that include an unlabeled pre-training phase, We first pre-train them for *N* epochs using the same unlabeled samples from both cancer cell lines and tumor tissues. With parameter grid search, *N* is selected based on the downstream task performance (over validation set). The pre-trained encoders will then be appended with a classification module to perform the downstream drug sensitivity classification task in the following fine-tuning step. We adopted the early stopping with validation performance in the fine-tuning phase (training phase for the model without unlabeled pre-training). Specifically, the labeled cell line samples were split into five stratified folds (according to drug sensitivity categorization). In one evaluation iteration, four out of five folds of the samples were used as the training set. The remaining one-fold of samples was used as the validation data set for early stopping. At last, the test performance of the classifier in each evaluation iteration was recorded.

#### 4.4.4 Performance evaluation

We choose the area under the receiver operating curve (AUROC) as the measurement metric due to their insensitivity to changes in the test data set’s class distribution [49]. The model performance was measured in AUROC over the patient tissue expression data and corresponding drug response records. The performance of different methods was compared by the average of AUROCs of five iterations. It is noted that only cell line data were used for the model training and hyperparameter selections, and all *ex vivo* tissues and patient data were purely used for the testing.

## Data availability

The original CCLE, GDSC, PDTC and TCGA data are publicly available datasets. CCLE data were downloaded from DepMap portal https://depmap.org/portal/download/. GDSC data were downloaded from the GDSC Website https://www.cancerrxgene.org/. PDTC datasets were obtained from Breast Cancer PDTX Encyclopaedia (https://caldaslab.cruk.cam.ac.uk/bcape/). TCGA data were downloaded from UCSC Cancer Genome Browser Xena [42]. Other intermediate files and TCGA tissue sample predictions can be found at https://github.com/XieResearchGroup/CODE-AE.

## Code availability

The source code and data are available at https://github.com/XieResearchGroup/CODE-AE.

## References

[1] Thai-Hoang Pham, Yue Qiu, Jucheng Zeng, Lei Xie, and Ping Zhang. A deep learning framework for high-throughput mechanism-driven phenotype compound screening and its application to covid-19 drug repurposing. Nature Machine Intelligence, 3(3):247–257, 2021.

[2] Jordi Barretina, Giordano Caponigro, Nicolas Stransky, Kavitha Venkatesan, Adam A Margolin, Sungjoon Kim, Christopher J Wilson, Joseph Lehár, Gregory V Kryukov, Dmitriy Sonkin, et al. The cancer cell line encyclopedia enables predictive modelling of anticancer drug sensitivity. Nature, 483(7391):603–607, 2012.

[3] Theodore Sakellaropoulos, Konstantinos Vougas, Sonali Narang, Filippos Koinis, Athanassios Kotsinas, Alexander Polyzos, Tyler J Moss, Sarina Piha-Paul, Hua Zhou, Eleni Kardala, et al. A deep learning framework for predicting response to therapy in cancer. Cell reports, 29(11):3367–3373, 2019.

[4] Ayse B Dincer, Joseph D Janizek, and Su-In Lee. Adversarial deconfounding autoencoder for learning robust gene expression embeddings. bioRxiv, 2020.

[5] Jianzhu Ma, Samson H Fong, Yunan Luo, Christopher J Bakkenist, John Paul Shen, Soufiane Mourragui, Lodewyk FA Wessels, Marc Hafner, Roded Sharan, Jian Peng, et al. Few-shot learning creates predictive models of drug response that translate from high-throughput screens to individual patients. Nature Cancer, 2(2):233–244, 2021.

[6] Peilin Jia, Ruifeng Hu, Guangsheng Pei, Yulin Dai, Yin-Ying Wang, and Zhongming Zhao. Deep generative neural network for accurate drug response imputation. Nature Communications, 12(1):1–16, 2021.

[7] Alejandra Bruna, Oscar M Rueda, Wendy Greenwood, Ankita Sati Batra, Maurizio Callari, Rajbir Nath Batra, Katherine Pogrebniak, Jose Sandoval, John W Cassidy, Ana Tufegdzic-Vidakovic, et al. A biobank of breast cancer explants with preserved intra-tumor heterogeneity to screen anticancer compounds. Cell, 167(1):260–274, 2016.

[8] Diederik P Kingma and Max Welling. Auto-encoding variational bayes. arXiv preprint arXiv:1312.6114, 2013.

[9] Pascal Vincent, Hugo Larochelle, Yoshua Bengio, and Pierre-Antoine Manzagol. Extracting and composing robust features with denoising autoencoders. In Proceedings of the 25th international conference on Machine learning, pages 1096–1103, 2008.

[10] Baochen Sun and Kate Saenko. Deep coral: Correlation alignment for deep domain adaptation. In European conference on computer vision, pages 443–450. Springer, 2016.

[11] Konstantinos Bousmalis, George Trigeorgis, Nathan Silberman, Dilip Krishnan, and Dumitru Erhan. Domain separation networks. In Advances in neural information processing systems, pages 343–351, 2016.

[12] Rotem Ben-Hamo, Alona Zilberberg, Helit Cohen, Keren Bahar-Shany, Chaim Wachtel, Jacob Korach, Sarit Aviel-Ronen, Iris Barshack, Danny Barash, Keren Levanon, et al. Resistance to paclitaxel is associated with a variant of the gene bcl2 in multiple tumor types. NPJ precision oncology, 3(1):1–11, 2019.

[13] Jenifer L Marks, Stephen Broderick, Qin Zhou, Dhananjay Chitale, Allan R Li, Maureen F Zakowski, Mark G Kris, Valerie W Rusch, Christopher G Azzoli, Venkatraman E Seshan, et al. Prognostic and therapeutic implications of egfr and kras mutations in resected lung adenocarcinoma. Journal of thoracic oncology, 3(2):111–116, 2008.

[14] Ben Markman, Francisco Javier Ramos, Jaume Capdevila, and Josep Tabernero. Egfr and kras in colorectal cancer. Advances in clinical chemistry, 51:72, 2010.

[15] Hui Hua, Qingbin Kong, Jie Yin, Jin Zhang, and Yangfu Jiang. Insulin-like growth factor receptor signaling in tumorigenesis and drug resistance: a challenge for cancer therapy. Journal of hematology & oncology, 13:1–17, 2020.

[16] Brian A Hemmings and David F Restuccia. Pi3k-pkb/akt pathway. Cold Spring Harbor perspectives in biology, 4(9):a011189, 2012.

[17] Michael S Isakoff, Courtney G Sansam, Pablo Tamayo, Aravind Subramanian, Julia A Evans, Christine M Fillmore, Xi Wang, Jaclyn A Biegel, Scott L Pomeroy, Jill P Mesirov, et al. Inactivation of the snf5 tumor suppressor stimulates cell cycle progression and cooperates with p53 loss in oncogenic transformation. Proceedings of the National Academy of Sciences, 102(49):17745–17750, 2005.

[18] Yong-Yeon Cho, Zhiwei He, Yiguo Zhang, Hong Seok Choi, Feng Zhu, Bu Young Choi, Bong Seok Kang, Wei-Ya Ma, Ann M Bode, and Zigang Dong. The p53 protein is a novel substrate of ribosomal s6 kinase 2 and a critical intermediary for ribosomal s6 kinase 2 and histone h3 interaction. Cancer research, 65(9):3596–3603, 2005.

[19] Yang Xu, Wensheng Yan, and Xinbin Chen. Snf5, a core component of the swi/snf complex, is necessary for p53 expression and cell survival, in part through eif4e. Oncogene, 29(28):4090–4100, 2010.

[20] Derek K Cheng, Tobiloba E Oni, Youngkyu Park, Jennifer S Thalappillil, Hsiu-chi Ting, Nadia Prasad, Brinda Alagesan, Keith D Rivera, Darryl J Pappin, Linda Van Aelst, et al. Oncogenic kras engages an rsk1/nf1 complex in pancreatic cancer. bioRxiv, 2020.

[21] Lixin Wan, Kexin Xu, Yongkun Wei, Jinfang Zhang, Tao Han, Christopher Fry, Zhao Zhang, Yao Vickie Wang, Liyu Huang, Min Yuan, et al. Phosphorylation of ezh2 by ampk suppresses prc2 methyltransferase activity and oncogenic function. Molecular cell, 69(2):279–291, 2018.

[22] Zonneville Justin, Vincent Wong, Limoge Michelle, Nikiforov Mikhail, and Andrei V Bakin. Tak1 signaling regulates p53 through a mechanism involving ribosomal stress. Scientific Reports (Nature Publisher Group), 10(1), 2020.

[23] Paul H Huang, Rebecca Cook, and Sibylle Mittnacht. Rb in dna repair. Oncotarget, 6(25):20746, 2015.

[24] Corey J Langer. Roles of egfr and kras mutations in the treatment of patients with non–small-cell lung cancer. Pharmacy and Therapeutics, 36(5):263, 2011.

[25] Inderjit S Dhillon. Co-clustering documents and words using bipartite spectral graph partitioning. In Proceedings of the seventh ACM SIGKDD international conference on Knowledge discovery and data mining, pages 269–274, 2001.

[26] David Arthur and Sergei Vassilvitskii. k-means++: The advantages of careful seeding. Technical report, Stanford, 2006.

[27] Kaisa Haglund, Tor Erik Rusten, and Harald Stenmark. Aberrant receptor signaling and trafficking as mechanisms in oncogenesis. Critical Reviews™ in Oncogenesis, 13(1), 2007.

[28] Charles Pottier, Margaux Fresnais, Marie Gilon, Guy Jérusalem, Rémi Longuespée, and Nor Eddine Sounni. Tyrosine kinase inhibitors in cancer: breakthrough and challenges of targeted therapy. Cancers, 12(3):731, 2020.

[29] Ana Jimenez-Pascual and Florian A Siebzehnrubl. Fibroblast growth factor receptor functions in glioblastoma. Cells, 8(7):715, 2019.

[30] Riccardo Riccardi, Devendra Singh, Joseph Minhow Chan, Pietro Zoppoli, Francesco Niola, Ryan Sullivan, Angelica Castano, Eric Minwei Liu, Jonathan Reichel, Paola Porrati, et al. Transforming fusions of fgfr and tacc genes in human glioblastoma. Science, pages N–A, 2012.

[31] Kathrin Schramm, Murat Iskar, Britta Statz, Natalie Jäger, Daniel Haag, Mikołaj Słabicki, Stefan M Pfister, Marc Zapatka, Jan Gronych, David TW Jones, et al. Decipher pooled shrna library screen identifies pp2a and fgfr signaling as potential therapeutic targets for diffuse intrinsic pontine gliomas. Neuro-oncology, 21(7):867–877, 2019.

[32] Han Kiat Ho, Sharon Pok, Sylvia Streit, Jens E Ruhe, Stefan Hart, Kah Suan Lim, Hooi Linn Loo, Myat Oo Aung, Seng Gee Lim, and Axel Ullrich. Fibroblast growth factor receptor 4 regulates proliferation, anti-apoptosis and alpha-fetoprotein secretion during hepatocellular carcinoma progression and represents a potential target for therapeutic intervention. Journal of hepatology, 50(1):118–127, 2009.

[33] Qing Wu, Aaron Bhole, Haiyan Qin, Judith Karp, Sami Malek, John K Cowell, and Mingqiang Ren. Targeting fgfr1 to suppress leukemogenesis in syndromic and de novo aml in murine models. Oncotarget, 7(31):49733, 2016.

[34] Di He and Lei Xie. A cross-level information transmission network for predicting phenotype from new genotype: Application to cancer precision medicine, 2020.

[35] Mathieu Salzmann, Carl Henrik Ek, Raquel Urtasun, and Trevor Darrell. Factorized orthogonal latent spaces. In Proceedings of the Thirteenth International Conference on Artificial Intelligence and Statistics, pages 701–708, 2010.

[36] Arthur Gretton, Karsten M Borgwardt, Malte J Rasch, Bernhard Schölkopf, and Alexander Smola. A kernel two-sample test. Journal of Machine Learning Research, 13(Mar):723–773, 2012.

[37] Martin Arjovsky, Soumith Chintala, and Léon Bottou. Wasserstein generative adversarial networks. In International conference on machine learning, pages 214–223. PMLR, 2017.

[38] Ishaan Gulrajani, Faruk Ahmed, Martin Arjovsky, Vincent Dumoulin, and Aaron C Courville. Improved training of wasserstein gans. In Advances in neural information processing systems, pages 5767–5777, 2017.

[39] Aravind Subramanian, Pablo Tamayo, Vamsi K Mootha, Sayan Mukherjee, Benjamin L Ebert, Michael A Gillette, Amanda Paulovich, Scott L Pomeroy, Todd R Golub, Eric S Lander, et al. Gene set enrichment analysis: a knowledge-based approach for interpreting genome-wide expression profiles. Proceedings of the National Academy of Sciences, 102(43):15545–15550, 2005.

[40] Vamsi K Mootha, Cecilia M Lindgren, Karl-Fredrik Eriksson, Aravind Subramanian, Smita Sihag, Joseph Lehar, Pere Puigserver, Emma Carlsson, Martin Ridderstraåle, Esa Laurila, et al. Pgc-1*α*-responsive genes involved in oxidative phosphorylation are coordinately downregulated in human diabetes. Nature genetics, 34(3):267–273, 2003.

[41] Mahmoud Ghandi, Franklin W Huang, Judit Jané-Valbuena, Gregory V Kryukov, Christopher C Lo, E Robert McDonald, Jordi Barretina, Ellen T Gelfand, Craig M Bielski, Haoxin Li, et al. Next-generation characterization of the cancer cell line encyclopedia. Nature, 569(7757):503–508, 2019.

[42] Mary Goldman, Brian Craft, Angela Brooks, Jing Zhu, and David Haussler. The ucsc xena platform for cancer genomics data visualization and interpretation. BioRxiv, page 326470, 2018.

[43] Wanjuan Yang, Jorge Soares, Patricia Greninger, Elena J Edelman, Howard Lightfoot, Simon Forbes, Nidhi Bindal, Dave Beare, James A Smith, I Richard Thompson, et al. Genomics of drug sensitivity in cancer (gdsc): a resource for therapeutic biomarker discovery in cancer cells. Nucleic acids research, 41(D1):D955–D961, 2012.

[44] Francesco Iorio, Theo A Knijnenburg, Daniel J Vis, Graham R Bignell, Michael P Menden, Michael Schubert, Nanne Aben, Emanuel Gonçalves, Syd Barthorpe, Howard Lightfoot, et al. A landscape of pharmacogenomic interactions in cancer. Cell, 166(3):740–754, 2016.

[45] Evan A Clayton, Toyya A Pujol, John F McDonald, and Peng Qiu. Leveraging tcga gene expression data to build predictive models for cancer drug response. BMC bioinformatics, 21(14):1–11, 2020.

[46] Carolyn Hutter and Jean Claude Zenklusen. The cancer genome atlas: creating lasting value beyond its data. Cell, 173(2):283–285, 2018.

[47] Geoffrey E Hinton and Richard S Zemel. Autoencoders, minimum description length, and helmholtz free energy. Advances in neural information processing systems, 6:3–10, 1994.

[48] Yaroslav Ganin, Evgeniya Ustinova, Hana Ajakan, Pascal Germain, Hugo Larochelle, François Laviolette, Mario Marchand, and Victor Lempitsky. Domain-adversarial training of neural networks. The Journal of Machine Learning Research, 17(1):2096–2030, 2016.

[49] Tom Fawcett. An introduction to roc analysis. Pattern recognition letters, 27(8):861–874, 2006.

