## Supplementary Materials for "Robust Prediction of Patient-Specific Clinical Response to Unseen Drugs From *in vitro* Screens Using Context-aware Deconfounding Autoencoder"

### A. Supplemental Table S1

**Table S1.** t-Test results on gene expression differences between responsive and resistant groups per drug-target pair

| Drug | Target | P-value | Avg. Expression (Responsive) | Avg. Expression (Resistant) |
|---|---|---|---|---|
| 681640 | WEE1 | 1.10E-22 | 4.0789 | 4.6994 |
| 681640 | CHEK1 | 7.90E-02 | 3.4623 | 3.324 |
| (5Z)-7-Oxozeaenol | MAP3K7 | 2.80E-07 | 3.4173 | 3.6632 |
| AG-014699 | PARP2 | 6.10E-05 | 4.2797 | 4.1018 |
| AG-014699 | PARP1 | 3.10E-01 | 6.0258 | 5.9679 |
| AICAR | PRKAA1 | 8.50E-101 | 3.5251 | 4.8481 |
| AMG-706 | PDGFRB | 6.90E-231 | 3.3645 | 6.0379 |
| AMG-706 | KDR | 2.20E-46 | 2.1994 | 3.2249 |
| AMG-706 | PDGFRA | 9.00E-20 | 2.9868 | 4.1865 |
| AMG-706 | RET | 1.10E-06 | 1.655 | 1.17 |
| AMG-706 | KIT | 7.90E-01 | 2.4167 | 2.3898 |
| Axitinib | PDGFRB | 4.00E-184 | 3.7331 | 6.1445 |
| Axitinib | PDGFRA | 5.80E-74 | 2.2025 | 4.2702 |
| Axitinib | KDR | 1.40E-24 | 2.5913 | 3.3782 |
| Axitinib | KIT | 4.50E-01 | 2.4023 | 2.4853 |
| AZ628 | BRAF | 2.40E-24 | 2.8583 | 3.2854 |
| AZD6482 | PIK3CB | 1.00E-03 | 3.8311 | 4.019 |
| AZD7762 | CHEK1 | 1.70E-185 | 4.1895 | 2.2045 |
| AZD7762 | CHEK2 | 5.40E-134 | 4.3098 | 2.6937 |
| AZD8055 | MTOR | 1.90E-01 | 4.245 | 4.1776 |
| BI-2536 | PLK2 | 2.60E-143 | 2.8942 | 5.4151 |
| BI-2536 | PLK3 | 2.90E-41 | 3.1475 | 3.8938 |
| BI-2536 | PLK1 | 4.40E-04 | 4.1665 | 3.8311 |
| BMS-536924 | IGF1R | 1.80E-72 | 2.3833 | 4.0068 |
| BMS-754807 | IGF1R | 9.60E-29 | 3.8051 | 2.9307 |
| Bosutinib | SRC | 2.30E-94 | 3.4436 | 4.8906 |
| Bosutinib | ABL1 | 4.80E-77 | 4.0001 | 5.1314 |
| Bosutinib | TEC | 1.10E-65 | 2.7255 | 1.1407 |
| BX-795 | AURKC | 4.80E-35 | 1.163 | 0.6489 |
| BX-795 | TBK1 | 3.70E-08 | 3.4926 | 3.822 |
| BX-795 | IKBKB | 2.20E-07 | 4.7648 | 4.4565 |
| BX-795 | AURKB | 3.50E-01 | 4.8741 | 4.7992 |
| BX-795 | PDK1 | 3.70E-01 | 2.8806 | 2.9369 |
| Camptothecin | TOP1 | 4.50E-111 | 4.4576 | 5.686 |
| CEP-701 | NTRK3 | 1.80E-40 | 2.3181 | 0.807 |
| CEP-701 | JAK2 | 3.10E-12 | 2.3605 | 2.8186 |

| | | | | |
|---|---|---|---|---|
| CEP-701 | NTRK2 | 7.10E-12 | 3.8394 | 2.4591 |
| CEP-701 | FLT3 | 1.10E-06 | 1.2452 | 0.7004 |
| CEP-701 | NTRK1 | 1.30E-05 | 0.8474 | 0.5553 |
| CHIR-99021 | GSK3A | 3.30E-03 | 5.9035 | 6.0019 |
| CHIR-99021 | GSK3B | 5.60E-01 | 4.1111 | 4.0846 |
| Embelin | XIAP | 5.00E-72 | 3.5752 | 4.2196 |
| FK866 | NAMPT | 3.60E-169 | 4.4288 | 6.6042 |
| GDC0941 | PIK3CA | 2.20E-69 | 2.7288 | 3.5768 |
| GDC0941 | PIK3CD | 9.00E-29 | 1.6778 | 2.2801 |
| Gefitinib | EGFR | 4.80E-47 | 3.9776 | 5.7095 |
| GSK1904529A | IGF1R | 2.90E-100 | 2.4047 | 3.9939 |
| GSK269962A | ROCK2 | 3.20E-20 | 3.2141 | 3.7583 |
| GSK269962A | ROCK1 | 2.10E-02 | 3.6288 | 3.469 |
| GW 441756 | NTRK1 | 1.50E-10 | 0.3287 | 0.6309 |
| JNJ-26854165 | MDM2 | 4.50E-07 | 5.7871 | 5.4236 |
| JNK Inhibitor VIII | MAPK8 | 2.50E-46 | 3.6722 | 4.5588 |
| JQ1 | BRD2 | 1.70E-08 | 6.9069 | 7.1648 |
| JQ1 | BRD4 | 2.70E-08 | 4.5332 | 4.7908 |
| JQ1 | BRDT | 4.00E-03 | 0.1288 | 0.255 |
| JQ1 | BRD3 | 1.90E-02 | 3.7328 | 3.8674 |
| KU-55933 | ATM | 3.00E-02 | 3.5916 | 3.4025 |
| LY317615 | PRKCB | 4.10E-10 | 3.6947 | 2.8402 |
| MK-2206 | AKT1 | 2.90E-106 | 5.5111 | 6.7304 |
| MK-2206 | AKT2 | 5.00E-39 | 5.5542 | 6.3624 |
| Nilotinib | ABL1 | 6.90E-87 | 3.7639 | 4.9984 |
| NU-7441 | PRKDC | 3.20E-01 | 5.5597 | 5.4917 |
| Obatoclax Mesylate | BCL2L2 | 1.50E-138 | 3.3805 | 4.8422 |
| Obatoclax Mesylate | BCL2L1 | 3.10E-88 | 5.5934 | 6.6164 |
| Obatoclax Mesylate | BCL2 | 1.60E-25 | 2.7184 | 3.7067 |
| Obatoclax Mesylate | MCL1 | 1.20E-15 | 6.8187 | 7.3133 |
| PAC-1 | CASP7 | 1.50E-173 | 3.3918 | 5.0627 |
| PAC-1 | CASP3 | 1.10E-70 | 3.709 | 4.6999 |
| PD-0325901 | MAP2K1 | 1.30E-17 | 4.5549 | 5.0474 |
| PD-0325901 | MAP2K2 | 5.70E-05 | 6.7388 | 6.9083 |
| PD-0332991 | CDK4 | 2.40E-43 | 6.4158 | 7.1613 |
| PD-0332991 | CDK6 | 2.60E-12 | 3.271 | 2.4058 |
| PD-173074 | FGFR1 | 1.80E-18 | 4.1757 | 4.9268 |
| PD-173074 | FGFR2 | 5.40E-11 | 3.1222 | 3.8407 |
| PD-173074 | FGFR3 | 3.10E-09 | 2.9842 | 3.7217 |
| PF-4708671 | RPS6KB1 | 2.10E-02 | 3.6593 | 3.5448 |
| PF-562271 | PTK2B | 3.10E-07 | 4.3494 | 4.0449 |
| PF-562271 | PTK2 | 8.50E-04 | 5.769 | 5.6216 |
| PLX4720 | BRAF | 1.80E-08 | 2.9273 | 3.2625 |
| RO-3306 | CDK1 | 2.30E-01 | 4.6398 | 4.737 |
| SB 216763 | GSK3A | 4.70E-44 | 5.4215 | 6.0307 |
| SB 216763 | GSK3B | 2.30E-11 | 3.7463 | 4.084 |
| SL 0101-1 | PIM3 | 5.10E-61 | 4.7294 | 5.7019 |
| SL 0101-1 | RPS6KA1 | 3.00E-11 | 4.5357 | 4.9659 |
| SL 0101-1 | PIM1 | 1.40E-05 | 4.7857 | 4.4367 |
| SL 0101-1 | AURKB | 4.10E-04 | 4.2786 | 3.9083 |
| Sorafenib | PDGFRB | 2.30E-222 | 2.6129 | 5.6893 |
| Sorafenib | PDGFRA | 1.20E-166 | 1.0217 | 3.7956 |
| Sorafenib | KDR | 5.70E-58 | 1.6047 | 2.8772 |
| Sorafenib | KIT | 2.10E-12 | 2.8686 | 2.0071 |

| | | | | |
|---|---|---|---|---|
| Sorafenib | RAF1 | 2.00E-10 | 5.0648 | 5.4119 |
| Tamoxifen | ESR1 | 2.20E-09 | 0.8943 | 1.5082 |
| TW 37 | MCL1 | 4.10E-106 | 6.7557 | 7.6602 |
| TW 37 | BCL2 | 3.80E-55 | 3.1667 | 1.9914 |
| TW 37 | BCL2L1 | 1.90E-17 | 6.3134 | 6.6994 |
| Vorinostat | HDAC1 | 1.90E-66 | 5.5303 | 6.6089 |
| Vorinostat | HDAC2 | 1.50E-44 | 5.1935 | 6.1185 |
| Vorinostat | HDAC6 | 2.70E-16 | 5.1075 | 4.7073 |
| Vorinostat | HDAC3 | 1.30E-15 | 5.1247 | 5.4736 |
| YK 4-279 | DHX9 | 1.20E-06 | 6.2476 | 6.4496 |
| ZM-447439 | AURKB | 3.00E-133 | 5.1265 | 2.8614 |
| ZM-447439 | AURKA | 5.10E-11 | 3.732 | 3.2247 |

### B. Supplemental Table S2

**Table S2.** TCGA tissue samples clustering results based on their drug responses towards 50 target drug therapies

| Drug | Tissue | Avg. Response Score | # of tissue samples / # of total samples within cluster |
|---|---|---|---|
| PD-173074 | brain lower grade glioma | 0.907573 | 351/372 |
| PD-173074 | pheochromocytoma & paraganglioma | 0.902569 | 105/214 |
| PD-173074 | liver hepatocellular carcinoma | 0.898942 | 197/382 |
| PD-173074 | acute myeloid leukemia | 0.883491 | 173/354 |
| PD-173074 | ovarian serous cystadenocarcinoma | 0.882003 | 55/190 |
| Camptothecin | liver hepatocellular carcinoma | 0.819653 | 84/289 |
| PD-0332991 | acute myeloid leukemia | 0.81628 | 173/354 |
| Camptothecin | acute myeloid leukemia | 0.814718 | 173/354 |
| Nilotinib | acute myeloid leukemia | 0.814684 | 173/354 |
| Camptothecin | pheochromocytoma & paraganglioma | 0.813179 | 105/214 |
| Camptothecin | brain lower grade glioma | 0.812126 | 351/372 |
| Vorinostat | acute myeloid leukemia | 0.80762 | 173/354 |
| Sorafenib | acute myeloid leukemia | 0.807489 | 173/354 |
| AICAR | head & neck squamous cell carcinoma | 0.804343 | 63/203 |
| Camptothecin | ovarian serous cystadenocarcinoma | 0.803714 | 55/190 |
| Sorafenib | pheochromocytoma & paraganglioma | 0.801101 | 105/214 |
| AICAR | acute myeloid leukemia | 0.795736 | 173/354 |
| Vorinostat | liver hepatocellular carcinoma | 0.795261 | 84/289 |
| Gefitinib | head & neck squamous cell carcinoma | 0.790417 | 63/203 |
| FK866 | brain lower grade glioma | 0.788069 | 351/372 |
| Sorafenib | liver hepatocellular carcinoma | 0.787842 | 84/289 |
| FK866 | liver hepatocellular carcinoma | 0.783162 | 197/382 |
| FK866 | pheochromocytoma & paraganglioma | 0.780261 | 105/214 |
| Vorinostat | ovarian serous cystadenocarcinoma | 0.780091 | 55/190 |
| FK866 | ovarian serous cystadenocarcinoma | 0.773476 | 55/190 |
| Sorafenib | brain lower grade glioma | 0.77254 | 351/372 |
| Tamoxifen | acute myeloid leukemia | 0.769454 | 173/354 |
| FK866 | acute myeloid leukemia | 0.767409 | 173/354 |
| Tamoxifen | ovarian serous cystadenocarcinoma | 0.748258 | 55/190 |
| PD-173074 | skin cutaneous melanoma | 0.747967 | 123/207 |
| Sorafenib | skin cutaneous melanoma | 0.74644 | 123/207 |
| PD-173074 | thyroid carcinoma | 0.744752 | 77/250 |
| PD-0332991 | liver hepatocellular carcinoma | 0.744704 | 84/289 |
| Gefitinib | acute myeloid leukemia | 0.74298 | 173/354 |
| AZD7762 | acute myeloid leukemia | 0.742207 | 173/354 |
| Vorinostat | pheochromocytoma & paraganglioma | 0.741767 | 105/214 |
| Sorafenib | testicular germ cell tumor | 0.741199 | 59/162 |
| Tamoxifen | liver hepatocellular carcinoma | 0.734867 | 84/289 |
| GSK1904529A | acute myeloid leukemia | 0.734821 | 173/354 |
| CHIR-99021 | sarcoma | 0.73268 | 183/352 |
| Nilotinib | liver hepatocellular carcinoma | 0.73167 | 84/289 |
| Embelin | head & neck squamous cell carcinoma | 0.725261 | 221/349 |
| Gefitinib | lung squamous cell carcinoma | 0.723799 | 204/471 |
| Sorafenib | glioblastoma multiforme | 0.722459 | 115/252 |
| AICAR | lung squamous cell carcinoma | 0.717723 | 204/471 |
| PD-173074 | testicular germ cell tumor | 0.717293 | 59/162 |
| PD-0325901 | thyroid carcinoma | 0.716933 | 145/225 |
| LY317615 | acute myeloid leukemia | 0.712712 | 173/354 |
| Nilotinib | testicular germ cell tumor | 0.711886 | 59/162 |
| YK 4-279 | acute myeloid leukemia | 0.711052 | 173/354 |
| Vorinostat | testicular germ cell tumor | 0.70557 | 59/162 |
| AICAR | testicular germ cell tumor | 0.704625 | 59/162 |
| Vorinostat | brain lower grade glioma | 0.701899 | 351/372 |
| PD-173074 | colon adenocarcinoma | 0.701831 | 45/149 |

### C. Supplemental Figure S1

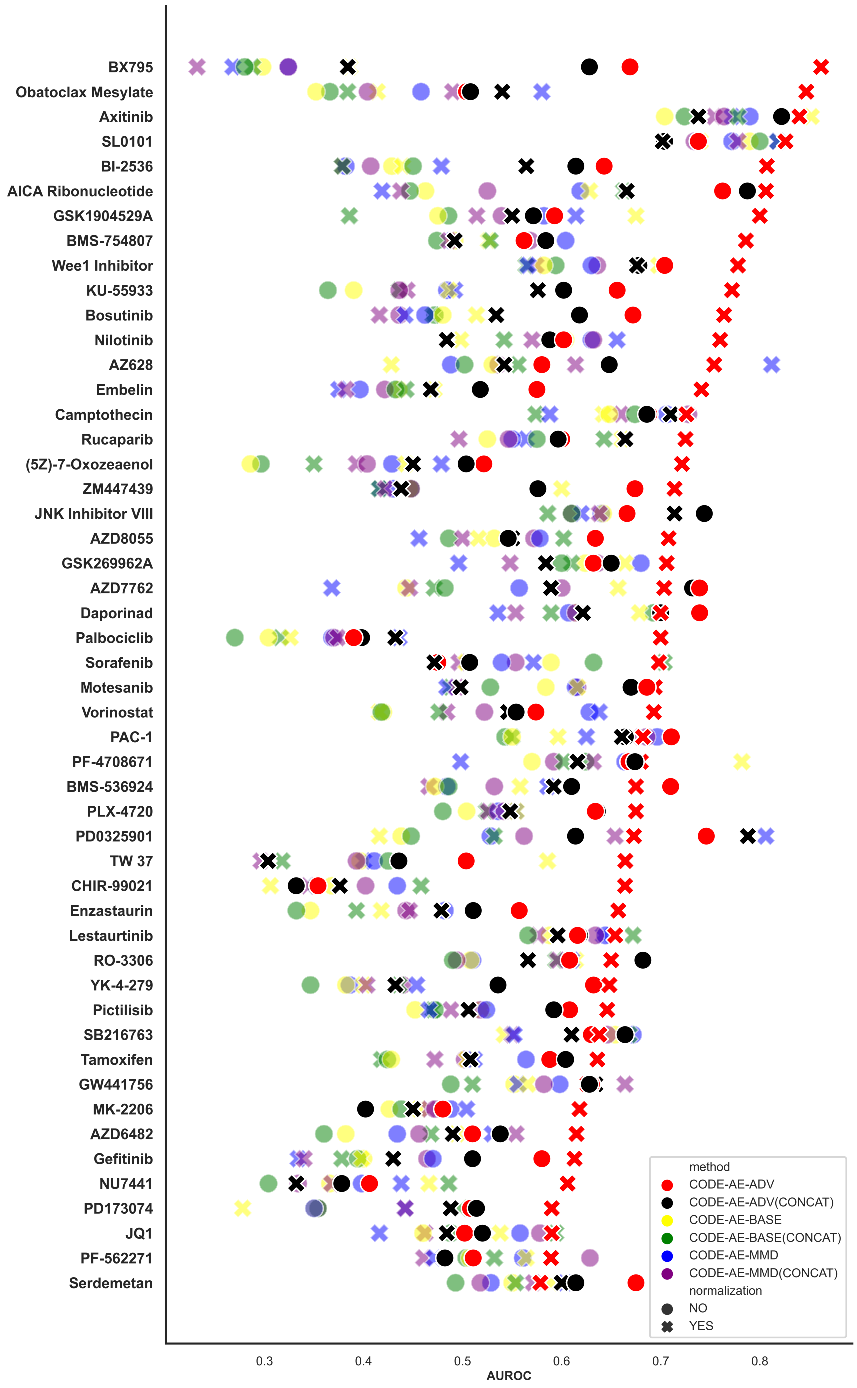


**Fig. S1.** PDTC drug response classification performance comparison of CODE-AE variants

### D. Supplemental Figure S2

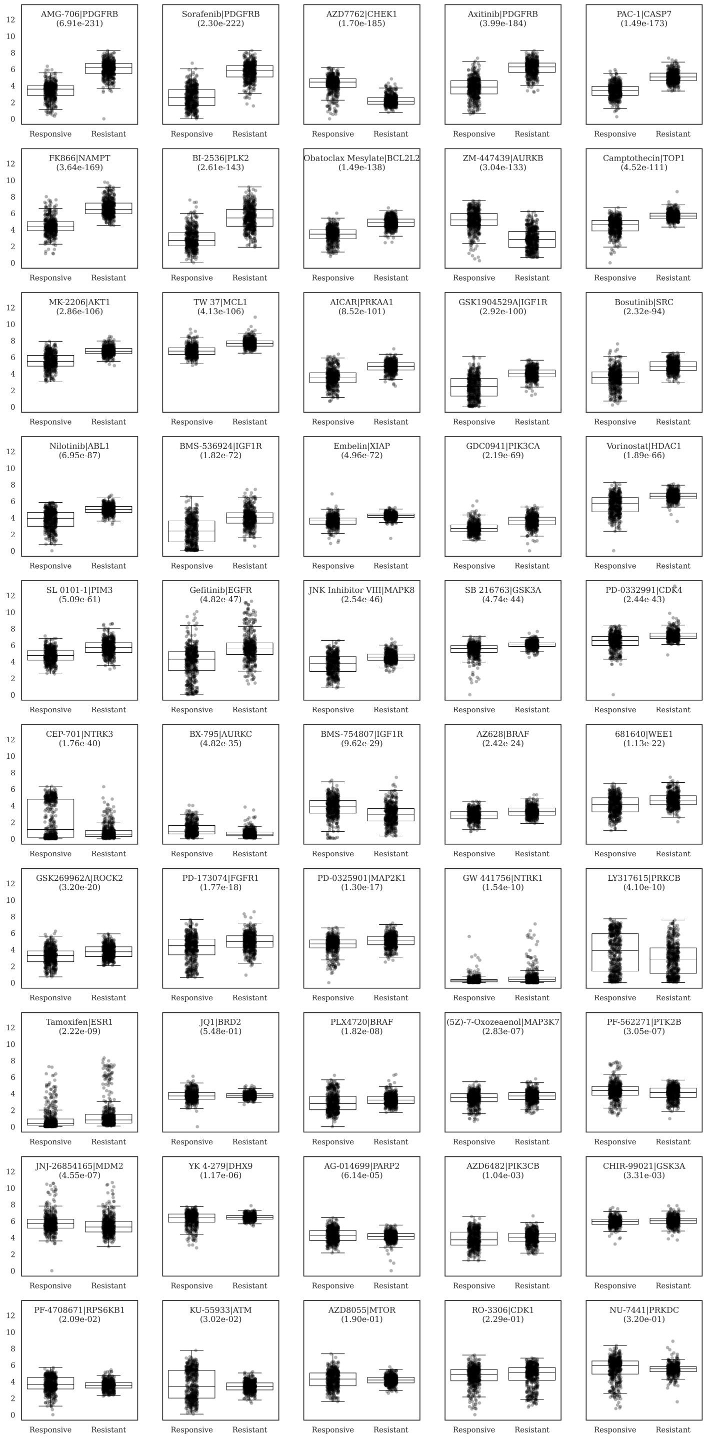


**Fig. S2.** Significant target gene expression value differences shown between responsive and resistant groups of 47 out of 50 target therapy agents

### E. Supplemental Figure S3

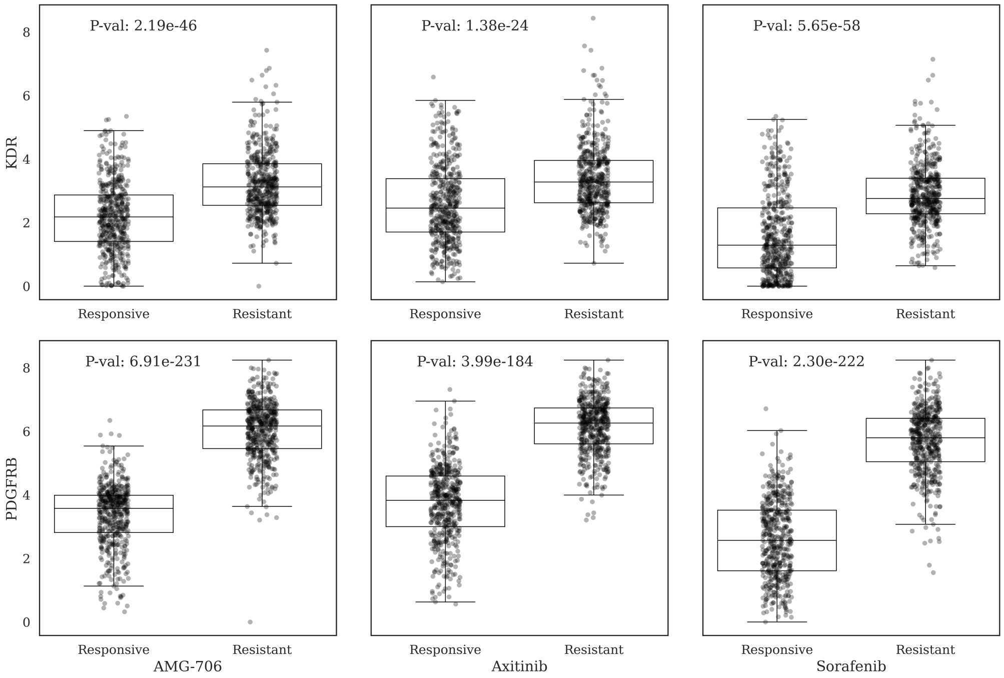


**Fig. S3.** Significant target gene (KDR and PDGFRB) expression value differences shown between responsive and resistant groups of drug Sorafenib, AMG-706 and Axitinib.
